# Truncated and “double-headed” derivatives of TO1-B dye for Mango-based imaging systems with increased brightness and selectivity and large Stocks shift

**DOI:** 10.1101/2024.12.11.627906

**Authors:** Julia I. Svetlova, Georgy K. Slushko, Anna S. Fayzieva, Evgeny S. Belyaev, Polina N. Kamzeeva, Andrey V. Aralov

**Author notes:** Equal contribution (Julia I. Svetlova, Georgy K. Slushko).

## Abstract

Fluorogenic dyes that light up in complexes with genetically encodable RNA aptamers are increasingly used for intracellular RNA imaging. The growing toolbox of dyes and aptamers allows simultaneous or sequential monitoring of different RNAs. However, few orthogonal aptamer-dye pairs meet all the requirements of multiplex imaging in terms of brightness, contrast, etc., thus necessitating their optimization and further diversification. Here, we report new fluorogenic ligands for the Mango II aptamer, one of the most stable and widely used RNA tags. They are based on the cognate Mango ligand TO1-B, which is a thiazole orange (TO) derivative with a biotinylated tetraethylene glycol (TEG) residue attached via an amide linker, and its analogue with an isosteric triazolyl linker (TO1-triazolyl-TEG-biotin). Structural data suggested that the biotin residue might be dispensable for interactions with Mango, so we replaced it with a second TO residue or the alternative push-pull system, namely the dimethylaminophenyl (DMAP) group linked to the benzothiazolyl (BzT) group via the ethenyl linker. The resulting “double-headed” symmetric and asymmetric dyes showed reduced fluorescence in the free state due to intramolecular TO/TO or TO/DMAP-BzT interactions. The asymmetric dye exhibited intramolecular FRET in complex with Mango, resulting in a remarkably large (130 nm) Stocks shift, which may be advantageous for multiplex imaging. The main limitation of the “double-headed” dyes was their low brightness. To improve brightness, we further optimized the truncated (biotin-free) TO1-triazolyl-TEG dye by introducing a hydroxy or a methoxy group into the methylqunolinium (MQ) fragment of its push-pull system. The brightness of the methoxy-MQ derivative was increased by 40% compared to the reported TO1-triazolyl-TEG-biotin. Importantly, the methoxy-MQ derivative also showed increased selectivity for Mango over other noncanonical nucleic acids structures, making it a promising alternative to known TO-based dyes for high-contrast RNA imaging.

## INTRODUCTION

Aptamers to fluorogenic dyes are valuable tools for intracellular RNA imaging ^[1,2]^. The aptamer module can be introduced into virtually any RNA as a genetically encoded tag to facilitate the detection of its localization, stimuli-driven redistribution, accumulation in specific cellular compartments or biocondensates, etc ^[3–6]^. Aptamer-arrays-based approaches are promising for single-molecule RNA monitoring^[7]^, and multiplex imaging techniques are opening the way to visualizing several different types of RNA in a given cell/cell culture^[8,9]^. The latter task requires a variety of aptamer-dye pairs with sufficient contrast and brightness, distinct spectral characteristics and minimal cross-reactivity. In this regard, much effort is devoted to the fine-tuning of the existing aptamers/dyes or the rational design of the new ones.

The current aptamer toolbox, including aptamers Spinach^[10]^, Broccoli^[11]^, Mango^[12]^, Corn^[13]^, Chili^[14]^, Pepper^[15]^, o-Coral^[16]^, Peach^[17]^, etc., and the possible orthogonal combinations with known dyes for multi-colour imaging are outlined in recent comprehensive reviews^[18]^. The majority of the aptamers adopt non-canonical secondary structures, such as G-quadruplexes (G4s)^[19–21]^. Stacks of planar guanine tetrads, together with G-tract-connecting loops and/or G-tract-interrupting bulges, provide facile structural platforms for interactions with conjugated aromatic systems of fluorogenic dyes. A classic example is the Mango family of aptamers (Mango I-IV ^[12,22]^) to cognate thiazole orange derivative (TO1-biotin) and related dyes. Each Mango aptamer contains a G/A-rich G4 core stabilized by a short duplex module. Crysal structures of Mango I-III complexes with TO derivatives show that loop residues enfold the external G-tetrad, creating a cavity that accommodates the TO structure and partially accommodates its substituents^[20,23–25]^. Within this cavity of Mango I aptamer, the conjugated fragments of the TO moiety, namely the 1-methylquinolinium (MQ) ring and the benzothiazolium (BzT) ring are stacked with G residues but rotated 45^°^ relative to each other leading to a moderate increase in the fluorescence quantum yield^[23]^. In contrast, the coplanarity of the fragments in complex with Mango II/III provokes a strong light-up effect^[20,24]^. Loop residues (e.g., A14, A25, A27, and A30 of Mango II) contribute to the overall stacking system and hold the dye in place (Fig. 1a).

**Fig. 1.**
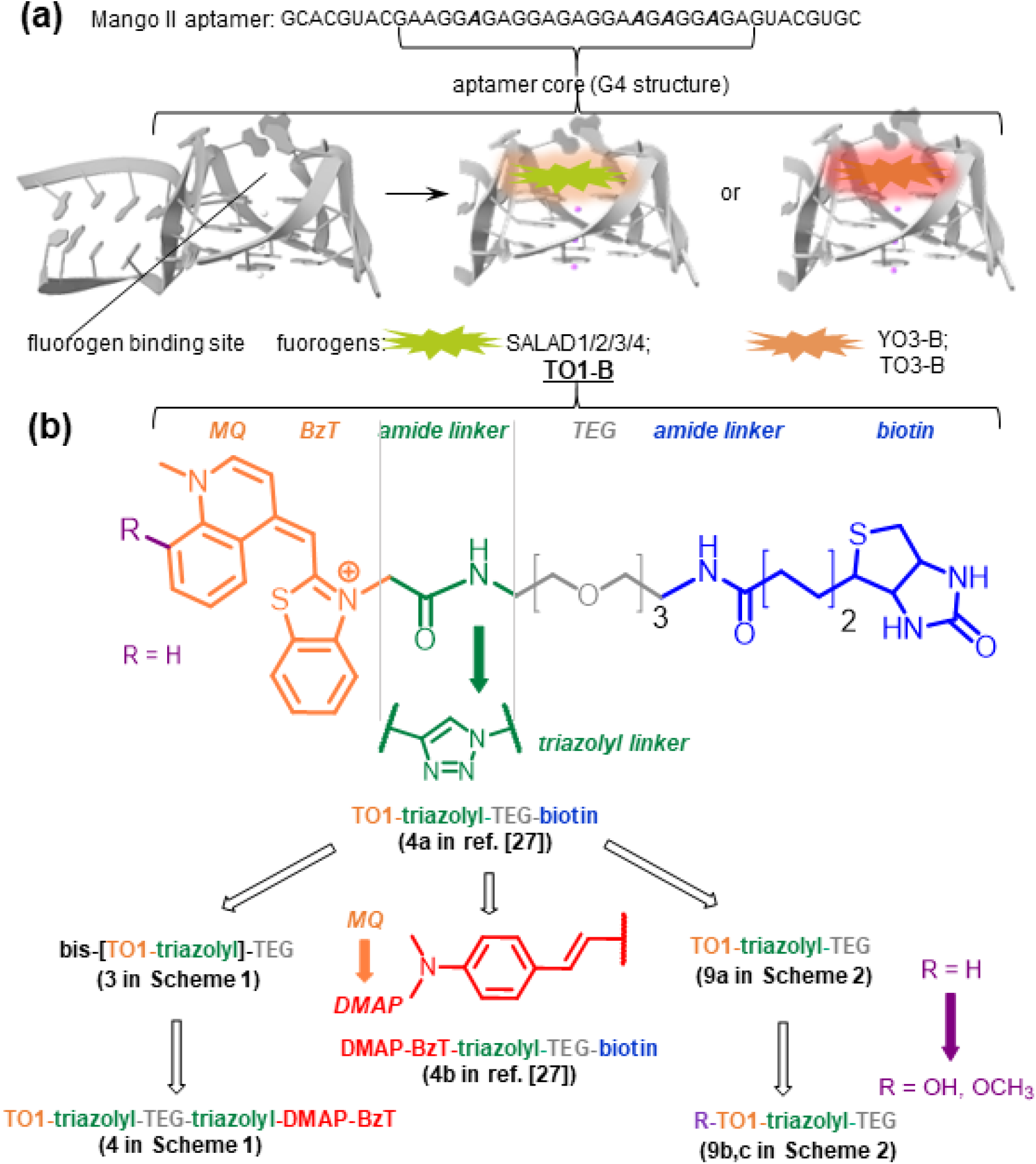
Schematic representation of the Mango-fluorogen light-up system (a) and the fluorogen design strategy (b).

Mango aptamers have been selected against a TO derivative conjugated to a biotinylated tetraethylene glycol (TEG) residue via an amide linker (TO1-B)^[22]^. The biotin-TEG fragment, although initially introduced for the selection step, proved advantageous in terms of the binding efficiency: the amide-TEG fragment occupied the TO-free space in the Mango binding site and contributed to complex stabilization, resulting in remarkably high binding affinities (Kd values in the low nanomolar range)^[20]^. In complex with Mango II, one of the most commonly used members of the aptamer family due to its thermal stability^[22]^, TO1 showed an approximately 1300-fold fluorescence light-up effect at 535 nm upon excitation at 510 nm^[20]^. While sufficient for proof-of-concept studies, this fluorescence properties of TO1-B needs to be further improved for intracellular applications and high-resolution RNA imaging.

Ongoing attempts to fine-tune the Mango-TO1-B pair aim to increase brightness/light-up, affinity and selectivity, or to modulate spectral parameters, including Stokes shift, for better compatibility with orthogonal pairs. Recently, a series of TO1-inspired Mango ligands have been developed by structure-informed design^[26]^. SALAD dyes with TO-like spectral properties but enhanced brightness and binding affinity for Mango II (Kd values in the sub-nanomolar range) were obtained by replacing the biotinylated TEG residue of TO1-B with a substituted benzyl ring. Beneficial substituents included the 1,2,3-triazole ring, which fitted into the Mango II cavity. In a previous work^[27]^, some of us also optimized TO1-B and analyzed the effects of replacing the amide linker between TO1 and TEG with the isosteric triazolyl linker (Fig. 1b). The target compound, TO1-triazolyl-TEG-biotin (**4a** in ^[27]^), was obtained by click ligation, and the increased yield at this ligation step was the main advantage of the modification. However, TO1-triazolyl-TEG-biotin also showed a slight increase in the light-up effect compared to TO1-B, probably due to the accommodation of the 1,2,3-triazole ring in the aptamer cavity.

Relevant TO1-B derivatives with altered spectral characteristics are those that emit in the red spectral region when bound to Mango. They can be used in multiplex imaging systems as orthogonal pairs for other green-light emitting dye-aptamer pairs or their FRET partners^[28]^, depending on the spectral overlap. Notable examples are the Mango I partners with extended conjugated systems, TO3 and YO3^[22,29]^. The former contains an extended trimethine linker between benzothiazole (BzT) and methylquinoline (MQ) residues of TO. The latter is an oxazole yellow derivative and a TO3 analogue with a benzoxazole ring instead of BzT. The spectra of both YO3 (excitation at 600 nm and emission at 630 nm) and TO3 (excitation at 620nm and emission at 670 nm) are shifted into the far-red region and show a modest overlap with the commonly used green-light emitting pairs DFHBI-Broccoli and DFHBI-1T-Spinach^[28]^, suggesting their applicability for orthogonal imaging but not for FRET-based imaging. Recently, we explored the possibility of tuning TO1-triazolyl-TEG-biotin by changing its push-and-pull (donor-π-acceptor) system and obtained a derivative with MQ substituted for the dimethylaminophenyl (DMAP) residue^[27]^. This derivative, DMAP-Bzt-triazolyl-TEG-biotin (**4b** in [27]) exhibited a moderate red shift and an increased Stokes shift (excitation at 560 nm and emission at 615 nm) in complex with Mango II, making it almost optimal for FRET-based systems with DFHBI-Broccoli. The remaining limitations of DMAP-Bzt-triazolyl-TEG-biotin and TO1-triazolyl-TEG-biotin are insufficient brightness and contrast.

In this work, we aimed to overcome the limitations of the above dyes and tested a new approach to contrast enhancement based on intramolecular quenching/FRET. We substituted the biotin moiety of TO1-triazolyl-TEG-biotin with a second fluorogenic moiety, TO1 or DMAP-BzT, yielding symmetric and asymmetric “double-headed” dyes, respectively. We also optimized the TO1-B derivative by introducing a hydroxy/methoxy group that can donate electrons through resonance into the donor (MQ) ring of its push-pull system and increase stacking interactions with surrounding nucleotide residues of the aptamer (Fig. 1b). Synthesis, spectral properties and selectivity of the new dyes are reported.

### Results and Discussion

### Synthesis and optical properties of symmetric and asymmetric “double-headed” dyes

The triazolyl-linked “double-headed” dyes **3** (bis-[TO1-triazolyl]-TEG) and **4** (TO1-triazolyl-TEG-triazolyl-DMAP-BzT) were synthesized following the reported procedure for copper catalyzed azide−alkyne cycloaddition with some modifications^[30]^ (**Scheme 1**). Briefly, TEG-bisazide **1** was condensed with **2** eq. of alkyne-containing fluorophore **2a** for the preparation of the symmetric dye **3**. The asymmetric dye **4** was synthesized by condensation of **2a** with a large excess of TEG-bisazide **1** (6 eq.), yielding an intermediate mono-azide, which was then condensed with **2b**. Control dyes with a single fluorogenic “head”, namely TO1-triazolyl-TEG-biotin (4a in [27]) and DMAP-BzT-triazolyl-TEG-biotin (**4b** in ref. [27]) were obtained as described previously^[27]^.

**Scheme 1.**
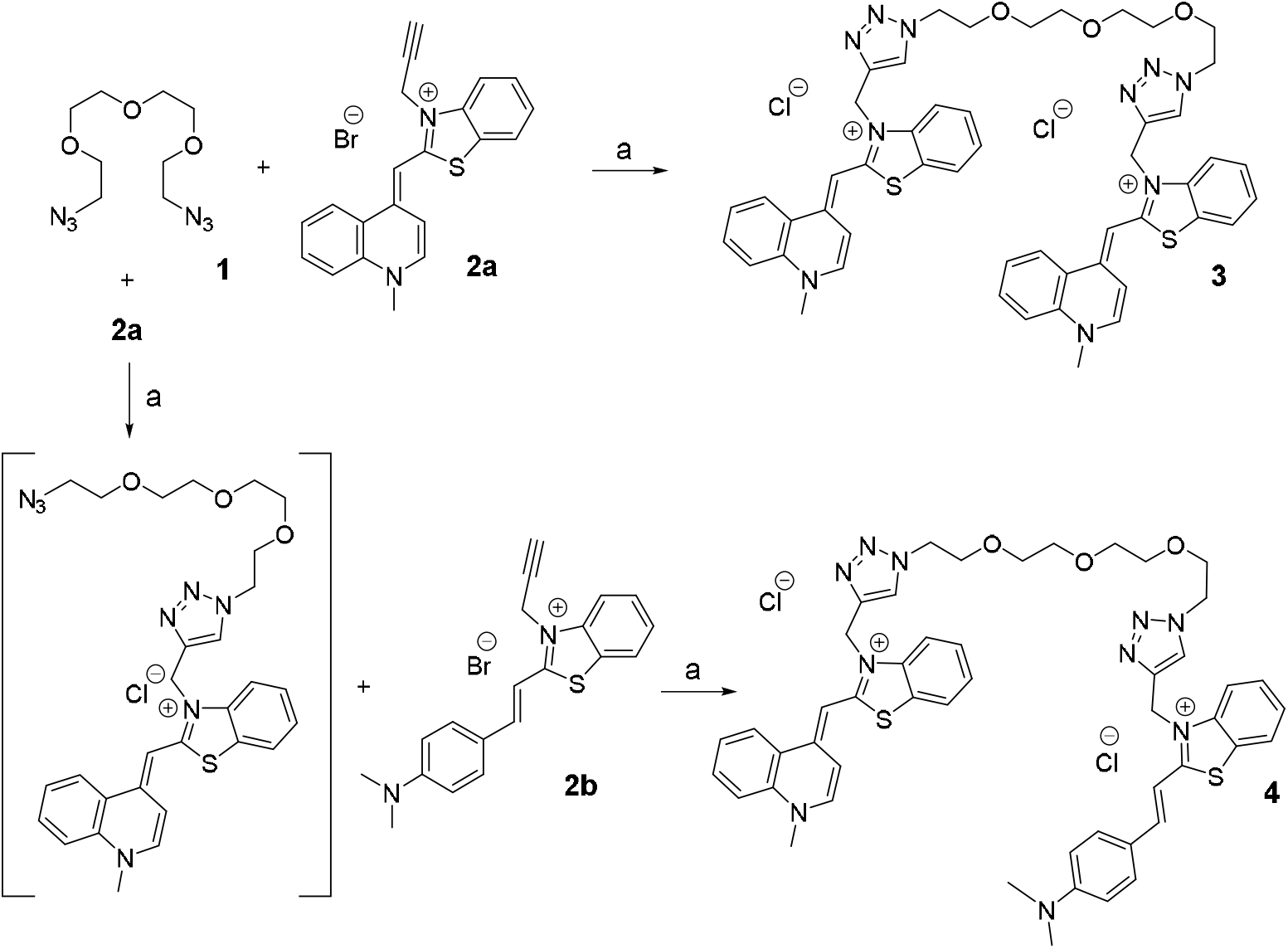
Synthesis of bis-[fluorophore-triazolyl]-TEG derivatives. Reagents and conditions: (a) CuI, TBTA, DMF, rt.

The optical properties of the free control (Fig. 2a) and new Fig. 2b) dyes in the pseudophysiological working buffer, as well as their complexes with Mango II, are summarized in Table 1. Both “double-headed” dyes showed absorption bands similar to those of TO1-triazolyl-TEG-biotin, except for a slight blue shift, indicating an excitonic interaction between the fluorogenic “heads”. Compound **4** had an additional broadened absorption band at 560-570 nm due to DMAP-BzT. The excitonic coupling of TO residues is known to suppress fluorescence emission and has previously been exploited to create DNA/RNA-binding light-up probes with improved contrast, such as TOTO homodimers^[31]^ or double-TO modified nucleoside analogues^[32]^.

**Table 1.** Spectral properties of the control and new dyes.

| dye | Free dye<br>Abs. max.<br>(nm) | Free dye $\epsilon$ at<br>Abs. max.<br>(M<sup>-1</sup>cm<sup>-1</sup>) | Mango-dye<br>$\lambda_{ex}$.<br>(nm) | Mango-dye<br>$\lambda_{em}$.<br>(nm) | $\Phi_{\text{Mango-dye}}$<br>(r.u.) | Brightness*<br>(M<sup>-1</sup>cm<sup>-1</sup>) |
| --- | --- | --- | --- | --- | --- | --- |
| TO1-triazolyl-TEG-biotin<br>(**4a** in ref. [27]) | 495 | 77500 | 505 | 540 | 0.42 | 29400 |
| DMAP-BzT-triazolyl-TEG-biotin (**4b** in ref.<br>[27]) | 535 | 77500 | 560 | 615 | 0.44 | 26200 |
| bis-[TO1-triazolyl]-TEG<br>(**3**) | 475 | 124000 | 480 | 550 | 0.01 | 1100 |
| TO1-triazolyl-TEG-triazolyl-DMAP-BzT (**4**) | 485 | 80000 | 505 | 615 (545) | 0.04 | 3200 |
| TO1-triazolyl-TEG<br>(**9a**) | 495 | 77500 | 510 | 540 | 0.28 | 26700 |
| HO-TO1-triazolyl-TEG<br>(**9b**) | 493 | 77500 | 510 | 545 | 0.40 | 37200 |
| CH<sub>3</sub>O-TO1-triazolyl-TEG<br>(**9c**) | 495 | 77500 | 510 | 540 | 0.45 | 40900 |

Brightness = $\Phi$ \*extinction at abs. max

**Fig. 2.**
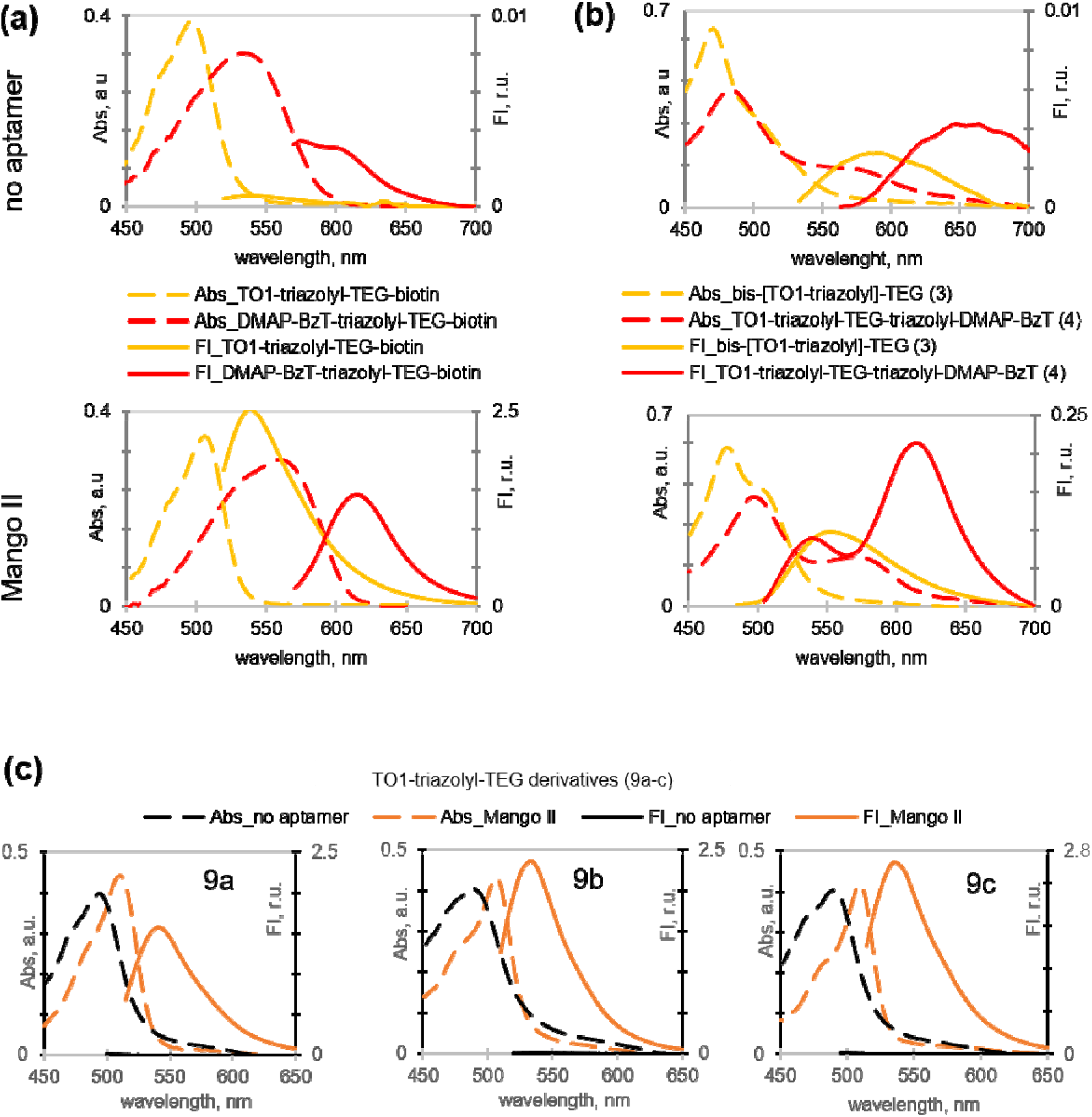
Spectral properties of the control and new dyes. Absorption and fluorescence emission spectra of control dyes (a), new “double-headed” dyes (b) and TO1-triazolyl-TEG derivatives (c) were obtained for free dye (5 µM solutions in the presudophysiological buffer) or their 1:1 mixtures with Mango II RNA.

For optimal contrast, the excitonic coupling between the fluorogens should be disrupted upon interaction with the aptamer. In the case of dye **3**, the disruption appears to be incomplete. In the presence of Mango II, dye **3** showed a shoulder absorption band at 500 nm, supporting the accommodation of at least one TO1 residue of **3** in the Mango cavity. However, the excitonic band at 475 nm was still present (Fig. 2b), suggesting that the second TO1 residue remained mostly stacked with the first one. As a result, the light-up effect of the symmetric dye **3** (approximately 40-fold fluorescence enhancement at 540-550 nm) was low compared to the control dye TO1-triazolyl-TEG-biotin (Table 2).

**Table 2.** Selectivity of the control and new dyes.

| Dye | Light-up* | $F_{\text{normalized}}^{**}$ | | |
| --- | --- | --- | --- | --- |
|  | Mangoll | 22AG | cKit-1 | Hairpin RNA |
| TO1-triazolyl-TEG-biotin (**4a** in ref. [27]) | 4770 | 14 | 15 | 11 |
| DMAP-BzT-triazolyl-TEG-biotin (**4b** in<br>ref. [27]) | 540 | 17 | 19 | 14 |
| bis-[TO1-triazolyl]-TEG (**3**) | 29 | 2 | 1,2 | 17 |
| TO1-triazolyl-TEG-triazolyl-DMAP-BzT<br>(**4**) | 8 | 30 | 11 | 6 |
| CH<sub>3</sub>O-TO1-triazolyl-TEG (**9c**) | 3150 | 661 | 33 | 11 |

The main absorption band of Mango-bound dye **4** coincided with that of TO1-triazolyl-TEG-biotin. Excitation at this band resulted in minor emission within the TO1-specific emission range (545 nm) and significant emission at the DMAP-BzT-specific wavelength (615 nm), which we attribute to intramolecular FRET (Fig. 2b). Thus, the TO1/DMAP-BzT stacking was altered or partially lost in complex with Mango, but FRET remained. The light-up effect of the asymmetric dye **4** at 615 nm was inferior to that of the control compound DMAP-BzT-triazolyl-TEG-biotin, but almost twice higher than that of the symmetric dye **3** at 560 nm (Table 2). We conclude that the asymmetric dye **4** is potentially useful for molecular imaging due to its remarkably large apparent Stokes shift (130 nm), resulting from intramolecular FRET between TO1 and DMAP-BzT moieties. The main disadvantage of dye **4** is its low brightness.

### Synthesis and optical properties of the biotin-free TO1-based dyes with a substituted MQ ring

The design of “double-headed” ligands proved beneficial in terms of the Stokes shift but did not allow us to improve the overall contrast. We therefore tried a different approach to optimizing TO1-triazolyl-TEG-biotin, focusing on the MQ ring. We explored the effects of hydroxy and methoxy substituents, which are too small to prevent the positioning of the modified TO1 residue in the Mango cavity but may alter its spectral properties due to the electron-donating resonance effect and provide additional contacts with the surrounding nucleotides of the aptamer. We also got rid of the biotin residue, which may be redundant, according to recent structural studies^[26]^. The synthesis of the biotin-free dyes with substituted MQ (TO1-triazolyl-TEG and its derivatives) is shown in **Scheme 2**.

**Scheme 2.**
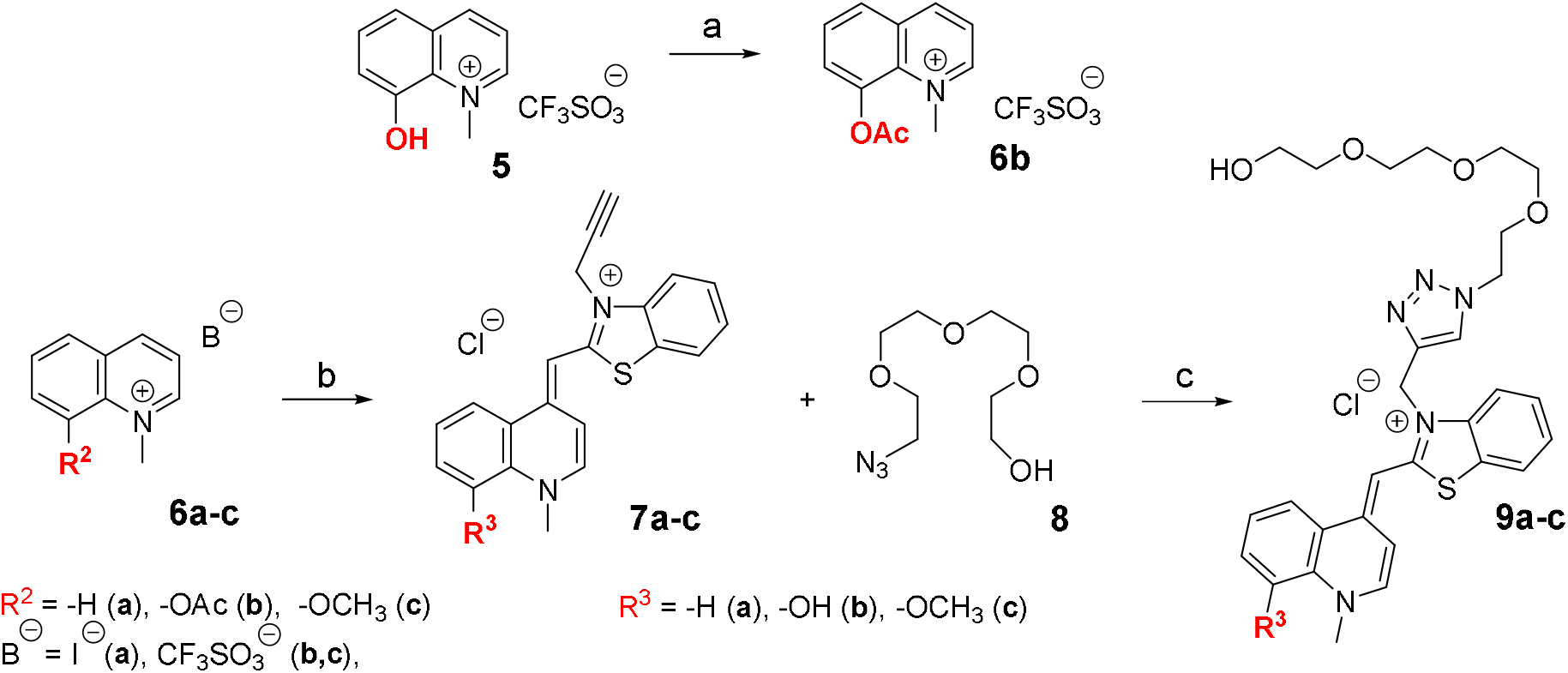
Synthesis of TO1-triazolyl-TEG derivatives. Reagents and conditions: (a) Ac_2_O, Py, rt; (b) 2-methyl-3-(prop-2-yn-1-yl)benzo[d]thiazol-3-ium bromide, TEA, CH_2_Cl_2_, rt; additional treatment with aqueous NH_3_ solution in CH_3_OH for 7b; (c) CuI, TBTA, DMF, rt.

The reported procedure of condensation between quinolinium salt **6a-c** and benzothiazolium salt under basic conditions was used to synthesize alkyne-containing TO1 derivatives **7a-c**^[12]^. To increase the condensation efficacy, the easily ionizable under basic conditions and thus electron-donating hydroxyl group of quinolinium salt **5** was protected with an acetyl group by the treatment with acetic anhydride in pyridine affording **6b**. Subsequent reaction of **7a-c** with TEG-monoazide **8** under «click reaction» conditions yielded target **9a-c**. Spectral properties of 9a-c are summarized in Table 1 and Fig. 2c.

The absorption and emission bands of **9a-c** were similar to those of TO1-triazolyl-TEG-biotin, except for the slightly increased Stokes shift. The fluorescence quantum yield of unsubstituted TO1-triazolyl-TEG (**9a**) was lower than that of TO1-triazolyl-TEG-biotin. Thus, although the biotin residue does not interact directly with the Mango cavity, it must contribute to the light-up effect – presumably, by stabilizing the complex. This is in line with the studies on amide-linked TO1-TEG derivatives^[20]^. The MQ substituents, especially the methoxy group, had a positive impact on the fluorescence quantum yield, and the leading new dye **9c** outperformed the control compound TO1-triazolyl-TEG-biotin. Due to the ease of synthesis and the increased brightness, this simplified ligand could be considered as a replacement for TO1-triazolyl-TEG-biotin and TO1-B in future studies.

### Analysis of the selectivity of the new dyes for Mango II over other structured nucleic acids

In addition to the pronounced light-up effect, high contrast intracellular imaging systems require the selectivity of the dye towards the aptamer over other nucleic acid structures, especially those prevalent in the cellular environment. Previously reported TO1 derivatives showed negligible cross-reactivity with presumably unstructured random-sequence ssRNA/DNA and minor to moderate cross-reactivity with structured DNA/RNA^[27]^. We focused on two types of nucleic acid structures: RNA hairpins, which are abundant in untranslated regions and other regulatory sites, and DNA G4s, which are abundant in genomic regulatory regions.

We used the following well-known sequences: ds26^[33]^ (Hairpin RNA), cKit-1 G4^[34]^ from c-KIT promotor, and 22AG G4 from telomeric repeats^[35]^. Secondary structures of all these sequences were confirmed by circular dichroism (CD) spectroscopy (Fig. 3a). Aptamer Mango II (a G4 with a short duplex module) and DNA G4s exhibited CD bands at 265 nm, consistent with the parallel-stranded quadruplex topology. DNA G4s showed an additional band at 295 nm. Thus, they have mixed topologies. The Mango II spectrum also contained a shoulder at 270-275 nm, which is a signature of the A-form duplex. The spectrum of Hairpin RNA was consistent with A-form with a positive band at 265 nm.

**Fig. 3.**
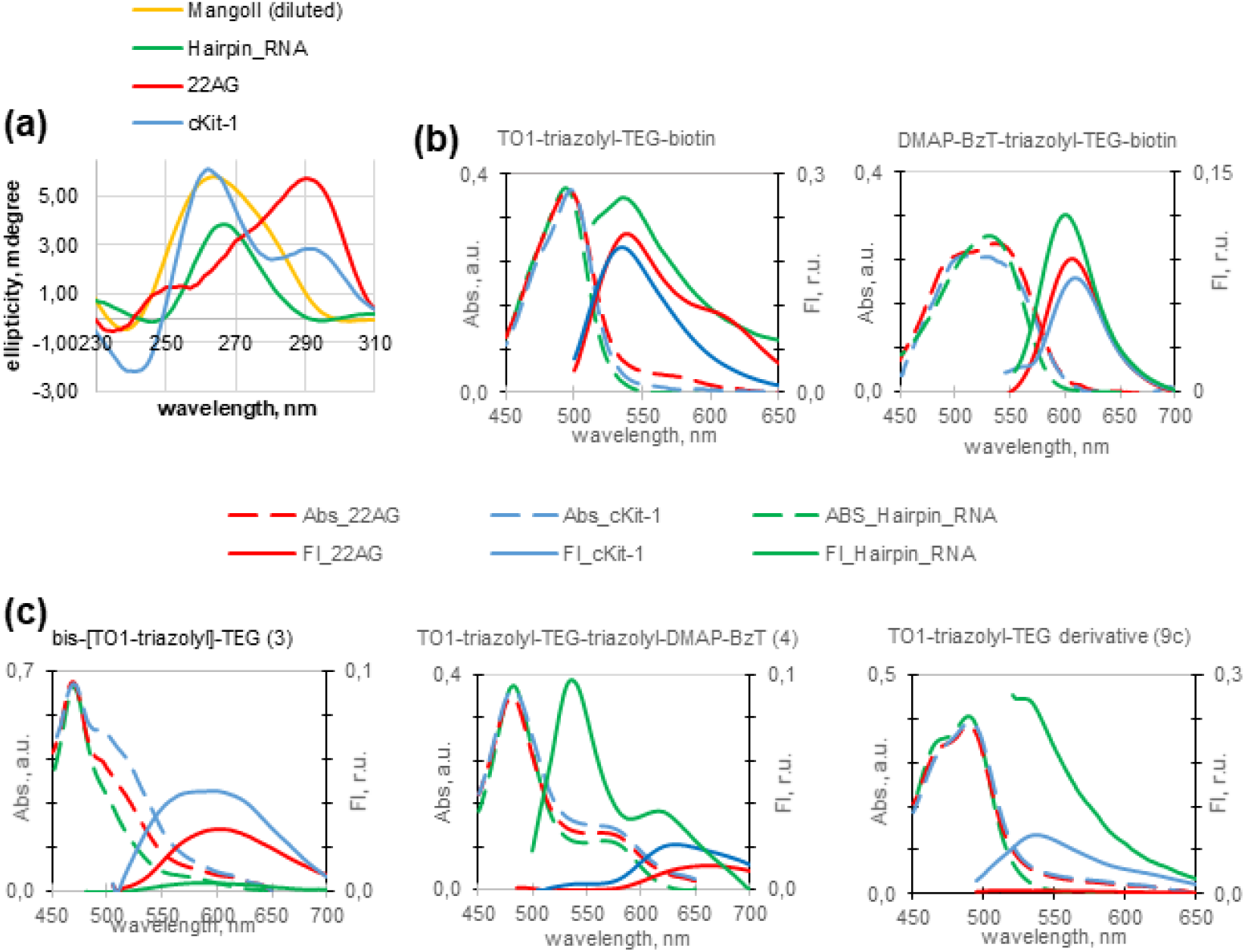
Spectral properties of the dyes in the presence of various RNA structures. (a) Circular dichroism spectra of Mango II (0.5 µM), Hairpin RNA (1 µM), 22AG (1 µM) and cKit-1 (1 µM) in the pseudophysiological buffer. (b) Absorption and fluorescence emission spectra of the control dyes (5 µM) in 1:1 mixture with Hairpin RNA/22AG/cKit-1. (c) Spectra of the new dyes (5 µM) in 1:1 mixture with Hairpin RNA/22AG/cKit-1.

Light-up effects of the “double-headed” dyes **3** and **4**, the control dyes TO1-triazolyl-TEG-biotin and DMAP-BzT-triazolyl-TEG-biotin, and the leading TO1-triazolyl-TEG derivative with the CH_3_O-substituted MQ ring (**9c**) were measured in the presence of the Hairpin RNA, 22AG, cKit-1 and compared to those in the presence of Mango II (Table 2). The control dyes showed moderate selectivity for Mango over the cKit-1, 22AG and Hairpin RNA (11-20-fold light-up difference). The symmetric “double-headed” dye **3** discriminated Mango from Hairpin but showed low selectivity over DNA G4s. The asymmetric dye **4** was inferior to **3** in terms of selectivity over Hairpin RNA. However, it outperformed dye **3** with cKit-1. It also outperformed **3** and the control dyes with 22AG. The new dye **9c** was similar to its control analogue TO1-triazolyl-TEG-biotin with Hairpin RNA and outperformed all tested dyes with DNA G4s. Its selectivity for Mango II over 22AG was remarkably high (660-fold light-up difference).

To summarize this part, improving selectivity for Mango over other G4s appears to be particularly challenging in the fluorogen design. Replacing TO1-triazolyl-TEG-biotin with new dyes **9c** or **4** can reduce off-target fluorescence, which is a significant step in the optimization of the Mango-based imaging systems.

## CONCLUSION

We designed new fluorogenic ligands for the Mango II aptamer, inspired by the TO1-B derivative with a triazolyl linkage (TO1-triazolyl-TEG-biotin) and its red-light emitting analogue (DMAP-BzT-triazolyl-TEG-biotin). The former was optimized by removing the biotin residue and introducing hydroxy/methoxy groups to the MQ ring. This resulted in a simplified synthetic protocol and afforded dyes **9b**,**c** with increased brightness in complex with Mango II. The methoxy-MQ dye **9c** showed increased selectivity for Mango over endogenous DNA G4s. In complex with Mango II, the new dye **9c** was 40% brighter than TO1-triazolyl-TEG-biotin and should be considered for high contrast intracellular imaging. We also designed a TO1-B derivative with an additional fluorogenic residue – TO1-triazolyl-TEG-triazolyl-DMAP-BzT **4**. This asymmetric dye was prone to intramolecular FRET and showed DMAP-BzT-specific far-red emission upon TO1 excitation in complex with Mango II. Despite the low quantum yield, this asymmetric dye could be used in multiplex imaging systems due to its relatively high selectivity and the remarkably large apparent Stokes shift (130 nm).

### Experimental section

#### Oligonucleotides, absorption spectroscopy, circular dichroism and fluorimetry

Sequences of all oligonucleotides (ONs) used in this study are listed below.

Mango II: r(GCACGUACGAAGGAGAGGAGAGGAAGAGGAGAGUACGUGC)

Hairpin RNA (da26): r(CAAUCGGAUCGAAUUCGAUCCGAUUG)

22AG: AGGGTTAGGGTTAGGGTTAGGG

cKit-1: AGGGAGGGCGCTGGGAGGAGGG

All ONs were obtained from GenTerra, Russia (>90% purity, HPLC). The 5 μM solutions in the pseudophysiological buffer (10 mM Tris–HCl, pH 7.5, and 140 mM KCl in MilliQ water) were annealed rapidly (heated to 95^°^C and cooled on ice) prior to the experiments. The dyes were added to preannealed ON solutions or blank solutions to a final concentration of 5 μM, affording 1:1 dye: ON mixtures. Absorption, circular dichroism and fluorescence emission spectra were registered with a Chirascan spectrophotometer (Applied Photophysics, UK) at room temperature in quartz cuvettes of 1 cm path.

Fluorescence quantum yields of the dyes were calculated using equation (1), TO1-B was used as a standard.

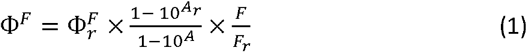

Where 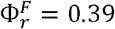 is fluorescence quantum yield of the reference dye TO1-B in the working buffer, *A*_r_ and *F*_r_ are absorbance and fluorescence of TO1-B in the working buffer, *A* and *F* are absorbance and fluorescence of the dye–RNA complex in the working buffer.

#### Synthesis of fluorogenic dyes

All reagents were commercially available unless otherwise mentioned and used without further purification. All solvents were purchased from commercial sources. Thin layer chromatography (TLC) was performed on plates (Merck) precoated with silica gel (60 mm, F254) and visualized using UV light (254 and 365 nm). Column chromatography (CC) was performed on silica gel (0.040-0.063 mm, Merck, Germany). ^1^H and ^13^C NMR spectra were recorded on the Bruker Avance III 600 spectrometer at 600 and 150 MHz, respectively. Chemical shifts are reported in δ (ppm) units using residual ^1^H signal from deuterated DMSO as reference. The multiplicity is reported using the following abbreviations: s (singlet), d (doublet), t (triplet), m (multiplet), br (broad). The coupling constants (J) are given in Hz. ESI HR mass spectra were acquired on a Thermo Scientific LTQ Orbitrap hybrid instrument (Thermo Electron Corp., Bremen, Germany) in continuous flow direct sample infusion (positive ion mode). 2-((1-Methylquinolin-4(1H)-ylidene)methyl)-3-(prop-2-yn-1-yl)benzo[d]thiazol-3-ium bromide **2a**^[27]^, 2-(2-(4-(dimethylamino)phenyl)ethenyl)-3-(2-propynyl)benzo[d]thiazolium bromide **2b**^[27]^, 8-hydroxy-1-methylquinolin-1-ium trifluoromethanesulfonate **5**^[36]^, 1-methylquinolinium iodide **6a**^[12]^, 8-methoxy-1-methylquinolinium trifluoromethanesulfonate **6c**^[37]^, and 2-methyl-3-(prop-2-yn-1-yl)benzo[d]thiazol-3-ium bromide^[38]^ were prepared according to the reported procedures. 1-Azido-2-(2-(2-(2-azidoethoxy)ethoxy)ethoxy)ethane 1 and 2-(2-(2-(2-azidoethoxy)ethoxy)ethoxy)ethan-1-ol **8** were purchased from Lumiprobe (Russia).

##### 3,3’-(((((oxybis(ethane-2,1-diyl))bis(oxy))bis(ethane-2,1-diyl))bis(1H-1,2,3-triazole-1,4-diyl))bis(methylene))bis(2-((-1-methylquinolin-4(1H)-ylidene)methyl)benzo[d]thiazol-3-ium) chloride 3

A solution of **1** (29 mg, 0.12 mmol) and **2a** (98 mg, 0.24 mmol, 2 eq) in DMF (10 mL) was degassed and then copper (I) iodide (4.6 mg, 0.024 mmol, 0.2 eq) and TBTA (6.4 mg, 0.012 mmol, 0.1 eq) were sequentially added under a nitrogen atmosphere. The reaction mixture was stirred at RT overnight, diluted with CH_2_Cl_2_ (20 mL) and washed with 1% aqueous EDTA solution (10 mL) and brine (10 mL). After concentration in vacuo, the residue was purified by column chromatography on silica gel (0-10% CH_3_OH in CH_2_ Cl_2_) yielding **3** (78 mg, 0.08 mmol, yield 67%) as red amorphous solid. ^1^H NMR (600 MHz, DMSO-d_6_) δ 8.62 – 8.50 (m, 4H, **H10, H10**’, **H6, H6**’), 8.34 – 8.25 (m, 2H, **H27, H27’**), 7.96 – 7.83 (m, 8H, **H8, H8**’, H12, **H12’, H18, H18’, H21, H21’**), 7.73 – 7.64 (m, 2H, **H9, H9’**), 7.56 – 7.47 (m, 2H, **H20, H20’**), 7.33 – 7.25 (m, 2H, **H19, H19’**), 7.22 – 7.13 (m, 4H, **H7, H7’, H5, H5’**), 5.90 – 5.78 (m, 4H, **H22, H22’**), 4.54 – 4.44 (m, 4H, **H28, H28’**), 4.13 (bs, 6H, **H11, H11’**), 3.77 – 3.67 (m, 4H, **H29, H29’**), 3.28 – 3.12 (m, 8H, **H31, H31’, H32, H32’**). ^13^C NMR (150 MHz, DMSO-d_6_) δ 158.5 (2C), 148.2 (2C), 144.8 (2C), 140.4 (2C), 139.3 (2C), 137.6 (2C), 132.8 (2C), 127.8 (2C), 126.7 (2C), 124.9 (2C), 124.3 (2C), 124.1 (2C), 123.8 (2C), 123.3 (2C), 122.5 (2C), 117.9 (2C), 112.6 (2C), 107.9 (2C), 88.2 (2C), 69.2 (2C), 69.1 (2C), 68.3(2C), 49.5 (2C), 42.3 (2C), 40.6 (2C). HRMS (ESI) m/z: calcd for C_50_ H_50_ N_10_ O_3_ S_2_ ^2+^ [M-2Cl^-^]^2+^: 451.1749; found 451.1748.

##### 2-(-4-(dimethylamino)styryl)-3-((1-(2-(2-(2-(2-(4-((2-((-1-methylquinolin-4(1H)-ylidene)methyl)benzo[d]thiazol-3-ium-3-yl)methyl)-1H-1,2,3-triazol-1-yl)ethoxy)ethoxy)ethoxy)ethyl)-1H-1,2,3-triazol-4-yl)methyl)benzo[d]thiazol-3-ium chloride 4

A solution of 1 (205 mg, 0.84 mmol) and **2a** (58 mg, 0.14 mmol) in DMF (20 mL) was degassed and then copper (I) iodide (2.7 mg, 0.014 mmol) and TBTA (3.7 mg, 0.007 mmol) were sequentially added under a nitrogen atmosphere. The reaction mixture was stirred at RT overnight and concentrated in vacuo. The residue was triturated with diethyl ether (3 x 20 mL) and dissolved in DMF (10 mL). To a resulting solution **2b** (56 mg, 0.14 mmol) was added followed by deaeration and addition of copper (I) iodide (2.7 mg, 0.014 mmol) and TBTA (3.7 mg, 0.007 mmol) under a nitrogen atmosphere. The reaction mixture was stirred at RT overnight, diluted with CH_2_Cl_2_ (20 mL) and washed with 1% aqueous EDTA solution (10 mL) and brine (10 mL). After concentration in vacuo, the residue was purified by column chromatography on silica gel (0-10% CH_3_OH in CH_2_Cl_2_) yielding **4** (82 mg, 0.09 mmol, yield 61% relative to **2a/2b**) as red amorphous solid. ^1^H NMR (600 MHz, DMSO-d_6_) δ 8.68 (d, J = 8.6 Hz, 1H, H63), 8.64 (d, J = 7.0 Hz, 1H, **H56**), 8.37 (s, 2H, **H17, H39**), 8.20 – 8.18 (m, 2H, **H35, H3**), 7.99 (d, J = 8.7 Hz, 1H, **H60**), 7.96 – 7.89 (m, 4H, **H25, H28, H61, H6**), 7.78 – 7.68 (m, 5H, **H62, H1, H51, H47, H34**), 7.61 (t, J = 7.7 Hz, 1H, **H2**), 7.56 (t, J = 7.8 Hz, 1H, **H24**), 7.35 (t, J = 7.6 Hz, 1H, **H23**), 7.31 – 7.28 (m, 2H, **H55, H12**), 6.70 (d, J = 8.8 Hz, 2H, **H50, H48**), 6.11 (s, 2H, **H10**), 5.89 (s, 2H, **H32**), 4.52 – 4.49 (m, 4H, **H18, H40**), 4.17 (s, 3H, **H64**), 4.13 – 4.11 (m, 2H, **H21**), 3.76 – 3.72 (m, 4H, **H19, H41**), 3.33 – 3.32 (m, 2H, overlaps with HDO, H43), 3.21 – 3.18 (m, 4H, **H44, H22**), 3.04 (s, 6H, **H53, H54**). HRMS (ESI) m/z: calcd for C_49_ H_45_ N_10_ O_3_ S_2_ ^2+^ [M-2Cl^-^]^2+^: 446.1827; found 446.1833.

##### 8-acetoxy-1-methylquinolin-1-ium trifluoromethanesulfonate 6b

To a solution of **5** (3.09 g, 10 mmol) in dry pyridine (20 mL) acetic anhydride (1.3 mL, 14 mmol) was added at RT. After stirring overnight at RT, CH_3_OH (1 mL) was added and the resulting mixture was concentrated in vacuo. The residue was triturated with diethyl ether (2 x 20 mL), acetone (5 mL) and dried in vacuo affording **6b** (3.02 g, 8.6 mmol, 86%) as brownish amorphous solid. ^1^H NMR (600 MHz, DMSO-d_6_) δ 9.47 (d, J = 5.7 Hz, 1H), 9.33 (d, J = 8.2 Hz, 1H), 8.40 (dd, J = 7.9, 1.7 Hz, 1H), 8.19 (dd, J = 8.4, 5.8 Hz, 1H), 8.11 (dd, J = 7.8, 1.7 Hz, 1H), 8.07 (t, J = 7.8 Hz, 1H), 4.72 (s, 3H), 2.53 (s, 3H). ^13^C NMR (150 MHz, DMSO-d_6_) δ 169.0, 153.2, 147.8, 146.9, 140.4, 131.5, 130.9, 129.8, 128.8, 122.3, 50.5, 21.4. HRMS (ESI) m/z: calcd for C_12_H_12_NO_2_^+^ [M-CF_3_SO_3_^-^]^+^: 202.0863; found 202.0862.

##### 2-((1-methylquinolin-4(1H)-ylidene)methyl)-3-(prop-2-yn-1-yl)benzo[d]thiazol-3-ium chloride 7a

This derivative was prepared from **6a** according to the reported procedure^[27]^ with additional washing with brine.

##### 2-((8-hydroxy-1-methylquinolin-4(1H)-ylidene)methyl)-3-(prop-2-yn-1-yl)benzo[d]thiazol-3-ium chloride 7b

To a suspension of **6b** (420 mg, 1.2 mmol) in CH_2_Cl_2_ (10 mL) 2-methyl-3-(prop-2-yn-1-yl)benzo[d]thiazol-3-ium bromide (320 mg, 1.2 mmol) and TEA (0.7 mL, 5 mmol) were added and the resulting mixture was stirred at RT overnight. Then the suspension was concentrated in vacuo and triturated with diethyl ether (2 x 10 mL). The residue was dissolved in CH_3_OH (40 mL) and aqueous ammonium hydroxide solution (1 mL) was added. After stirring overnight at RT, the reaction mixture was concentrated in vacuo, dissolved in a mixture of CH_3_OH/CH_2_Cl_2_ (1:20, 30 mL) and washed with brine (30 mL) in a separatory funnel. The solid formed was filtered, washed with water (2 x 10 mL) and dried in vacuo affording **7b** (88 mg, 0.23 mmol, 19%) as red amorphous solid. ^1^H NMR (600 MHz, DMSO-d_6_) δ 8.23 (d, J = 6.7 Hz, 1H), 7.84 (d, J = 7.8 Hz, 1H), 7.57 (d, J = 8.2 Hz, 1H), 7.48 (t, J = 7.9 Hz, 1H), 7.29 – 7.24 (m, 3H), 7.02 (d, J = 8.2 Hz, 1H), 6.66 (d, J = 8.1 Hz, 1H), 6.61 (s, 1H), 5.29 (s, 2H), 4.71 (s, 3H), 3.47 (s, 1H). ^13^C NMR (150 MHz, DMSO-d_6_) δ 154.1, 148.9, 143.6, 143.5, 139.6, 131.7, 129.4, 128.7, 127.6, 123.4, 122.8, 122.6 (2C), 119.8, 111.3, 109.4, 88.0, 77.1, 75.9, 48.4, 34.7. HRMS (ESI) m/z: calcd for C _21_ H_17_ N_2_ OS^+^ [M-Cl^-^]^+^: 345.1056; found 345.1056.

##### 2-((8-methoxy-1-methylquinolin-4(1H)-ylidene)methyl)-3-(prop-2-yn-1-yl)benzo[d]thiazol-3-ium chloride 7c

To a suspension of **6c** (390 mg, 1.2 mmol) in CH_2_Cl_2_ (10 mL) 2-methyl-3-(prop-2-yn-1-yl)benzo[d]thiazol-3-ium bromide (320 mg, 1.2 mmol) and TEA (0.7 mL, 5 mmol) were added and the resulting mixture was stirred at RT overnight. Then the suspension was concentrated in vacuo and triturated with diethyl ether (2 x 10 mL). The residue was dissolved in a mixture of CH_3_OH/CH_2_Cl_2_ (1:20, 30 mL) and washed with brine (30 mL) in a separatory funnel. The solid formed was filtered, washed with water (2 x 10 mL) and dried in vacuo affording **7c** (90 mg, 0.25 mmol, 21%) as red amorphous solid. ^1^H NMR (600 MHz, DMSO-d_6_) δ 8.58 (d, J = 7.2 Hz, 1H), 8.27 (d, J = 8.9 Hz, 1H), 8.01 (d, J = 7.9 Hz, 1H), 7.79 – 7.71 (m, 2H), 7.63 – 7.57 (m, 2H), 7.42 – 7.38 (m, 2H), 6.90 (s, 1H), 5.53 (s, 2H), 4.41 (s, 3H), 4.02 (s, 3H), 3.56 – 3.51 (m, 1H). ^13^C NMR (150 MHz, DMSO-d_6_) δ 158.1, 151.3, 148.6, 147.5, 139.2, 130.0, 128.2, 127.7, 126.6, 124.5, 123.4, 123.0, 116.9, 115.1, 112.4, 109.2, 88.4, 76.8, 76.4, 57.0, 49.4, 35.4. HRMS (ESI) m/z: calcd for C _22_H_19_N_2_OS^+^ [M-Cl^-^]^+^: 359.1213; found 359.1211.

#### General method for the preparation of TO-triazolyl-TEG derivatives 9a-c

A solution of **7** (0.12 mmol) and **8** (26 mg, 0.12 mmol, 1 eq) in DMF (10 mL) was degassed and then copper (I) iodide (2.3 mg, 0.012 mmol, 0.1 eq) and TBTA (3.2 mg, 0.006 mmol, 0.05 eq) were sequentially added under a nitrogen atmosphere. The reaction mixture was stirred at RT overnight, diluted with CH_2_Cl_2_ (20 mL) and washed with 1% aqueous EDTA solution (10 mL) and brine (10 mL). After concentration in vacuo, the residue was purified by column chromatography on silica gel (0-10% CH_3_OH in CH_2_Cl_2_) yielding **9** as red amorphous solid:

##### 3-((1-(2-(2-(2-(2-hydroxyethoxy)ethoxy)ethoxy)ethyl)-1H-1,2,3-triazol-4-yl)methyl)-2-((1-methylquinolin-4(1H)-ylidene)methyl)benzo[d]thiazol-3-ium chloride 9a

Yield 88%. ^1^H NMR (600 MHz, DMSO-d_6_) δ 8.78 (d, J = 7.9 Hz, 1H, **H35**), 8.70 (d, J = 7.3 Hz, 1H, **H33**), 8.36 – 8.33 (m, 1H, **H28**), 8.10 – 8.05 (m, 1H, **H11**), 8.05 – 7.98 (m, 2H, **H37, H6**), 7.98 – 7.93 (m, 1H, **H3**), 7.85 – 7.79 (m, 1H, **H36**), 7.63 – 7.57 (m, 1H, **H2**), 7.41 – 7.31 (m, 3H, **H1, H38, H34**), 5.92 (s, 2H, **H10**), 4.61 (s, 1H, **H23**), 4.51 (d, J = 4.9 Hz, 2H, **H12**), 4.19 (s, 3H, **H39**), 3.80 – 3.75 (m, 2H, **H13**), 3.48 – 3.33 (m, 12H, overlaps with HDO, H15-**H22**). ^13^C NMR (150 MHz, DMSO-d_6_) δ 158.9, 148.7, 145.3, 140.5, 139.8, 138.0, 133.2, 128.1, 127.1, 125.4, 124.6, 124.4, 124.2, 123.6, 122.9, 118.3, 113.0, 108.2, 88.5, 72.2, 69.7, 69.6, 69.5, 69.4, 68.6, 60.1, 49.5, 42.5, 40.9. HRMS (ESI) m/z: calcd for C_29_ H_34_N_5_O_4_S^+^ [M-Cl^-^]^+^: 548.2326; found 548.2324.

##### 2-((8-hydroxy-1-methylquinolin-4(1H)-ylidene)methyl)-3-((1-(2-(2-(2-(2-hydroxyethoxy)ethoxy)ethoxy)ethyl)-1H-1,2,3-triazol-4-yl)methyl)benzo[d]thiazol-3-ium chloride 9b

Yield 88%. ^1^H NMR (600 MHz, DMSO-d_6_) δ 11.24 (br s, 1H, **H40**), 8.49 (d, J = 7.1 Hz, 1H, **H33**), 8.30 (s, 1H, **H28**), 8.11 (d, J = 8.3 Hz, 1H, **H35**), 7.98 (d, J = 7.8 Hz, 1H, **H3**), 7.91 (d, J = 8.3 Hz, 1H, **H37**), 7.62 – 7.56 (m, 2H, **H36, H1**), 7.43 (d, J = 7.8 Hz, 1H, **H6**), 7.38 (t, J = 7.6 Hz, 1H, **H2**), 7.31 (d, J = 7.0 Hz, 1H, **H34**), 7.16 (s, 1H, **H11**), 5.84 (s, 2H, **H10**), 4.55 (br s, 1H, **H23**), 4.51 (t, J = 5.1 Hz, 2H, **H12**), 4.44 (s, 3H, **H39**), 3.77 (t, J = 5.1 Hz, 2H, **H13**), 3.49 – 3.36 (m, 12H, **H15-H22**, overlaps with HDO). ^13^C NMR (150 MHz, DMSO-d_6_) δ 158.2, 149.7, 148.6, 146.8, 140.5, 139.8, 128.8, 128.0, 127.7, 126.9, 124.5, 124.2, 123.4, 122.8, 118.4, 115.4, 112.7, 108.4, 88.6, 72.2, 69.6, 69.6, 69.5, 69.4, 68.6, 60.1, 49.5, 48.8, 40.8. HRMS (ESI) m/z: calcd for C_29_H_34_N_5_O_5_S^+^ [M-Cl^-^]^+^: 564.2275; found 564.2273.

##### 3-((1-(2-(2-(2-(2-hydroxyethoxy)ethoxy)ethoxy)ethyl)-1H-1,2,3-triazol-4-yl)methyl)-2-((8-methoxy-1-methylquinolin-4(1H)-ylidene)methyl)benzo[d]thiazol-3-ium chloride 9c

Yield 85%. ^1^H NMR (600 MHz, DMSO-d _6_) δ 8.47 (d, J = 7.1 Hz, 1H, **H33**), 8.31 (s, 1H, **H28**), 8.24 (d, J = 8.5 Hz, 1H, **H35**), 7.97 (d, J = 7.8 Hz, 1H, **H3**), 7.92 (d, J = 8.3 Hz, 1H, **H6**), 7.70 (t, J = 8.2 Hz, 1H, **H36**), 7.62 – 7.57 (m, 1H, **H1**), 7.55 (d, J = 7.8 Hz, 1H, **H37**), 7.38 (t, J = 7.6 Hz, 1H, **H2**), 7.30 (d, J = 7.1 Hz, 1H, **H34**), 7.18 (s, 1H, **H11**), 5.86 (s, 2H, **H10**), 4.56 – 4.49 (m, 3H, **H23, H12**), 4.36 (s, 3H, **H39**), 4.00 (s, 3H, **H40**), 3.78 (t, J = 5.1 Hz, 2H, **H22**), 3.46 – 3.34 (m, 12H, **H13-H21**, overlaps with HDO). ^13^C NMR (150 MHz, DMSO-d_6_) δ 158.7, 151.2, 148.3, 147.1, 140.5, 139.7, 129.9, 128.0, 127.5, 126.5, 124.5, 124.3, 123.5, 122.8, 116.9, 114.9, 112.8, 108.6, 88.8, 72.2, 69.6, 69.6, 69.5, 69.4, 68.6, 60.1, 56.9, 49.5, 49.1, 40.9. HRMS (ESI) m/z: calcd for C_30_H_36_N_5_O_5_S^+^ [M-Cl^-^]^+^: 578.2432; found 578.2431.

## Supporting information

Supplemenary file with NMR spectra

## Author contributions

Conceptualization of study and methodology (AVA); investigation (JIS, GKS, ASF, ESB, PNK); visualization and writing of original draft (JIS, GKS, AVA); writing — review and editing (AVA); funding acquisition (JIS), project administration, and supervision (AVA).

## Data availability

The data supporting this article have been included as part of the ESI.†

## Acknowledgements

This work has been supported by the grants the Russian Science Foundation, RSF 24-25-00486.

