## Supplementary material for "Truncated and “double-headed” derivatives of TO1-B dye for Mango-based imaging systems with increased brightness and selectivity and large Stocks shift": Supplemenary file with NMR spectra

### **Figures and Tables**

#### **Fig. S1. ^1^H spectrum of 3**

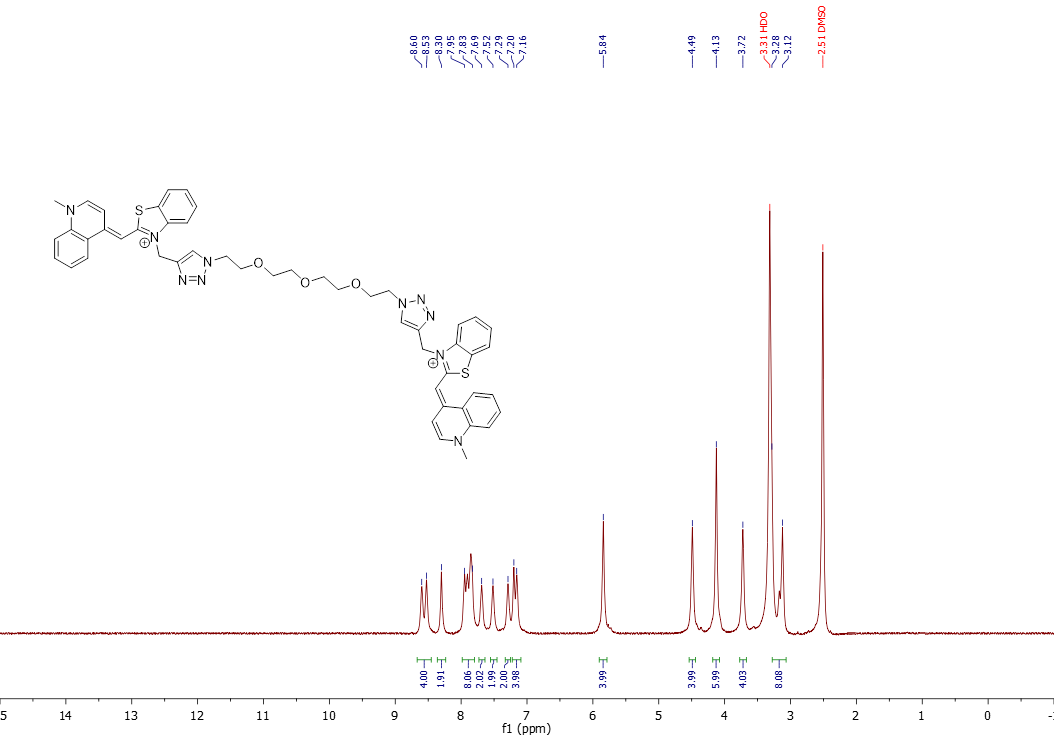

#### **Fig. S2. ^13^C spectrum of 3**

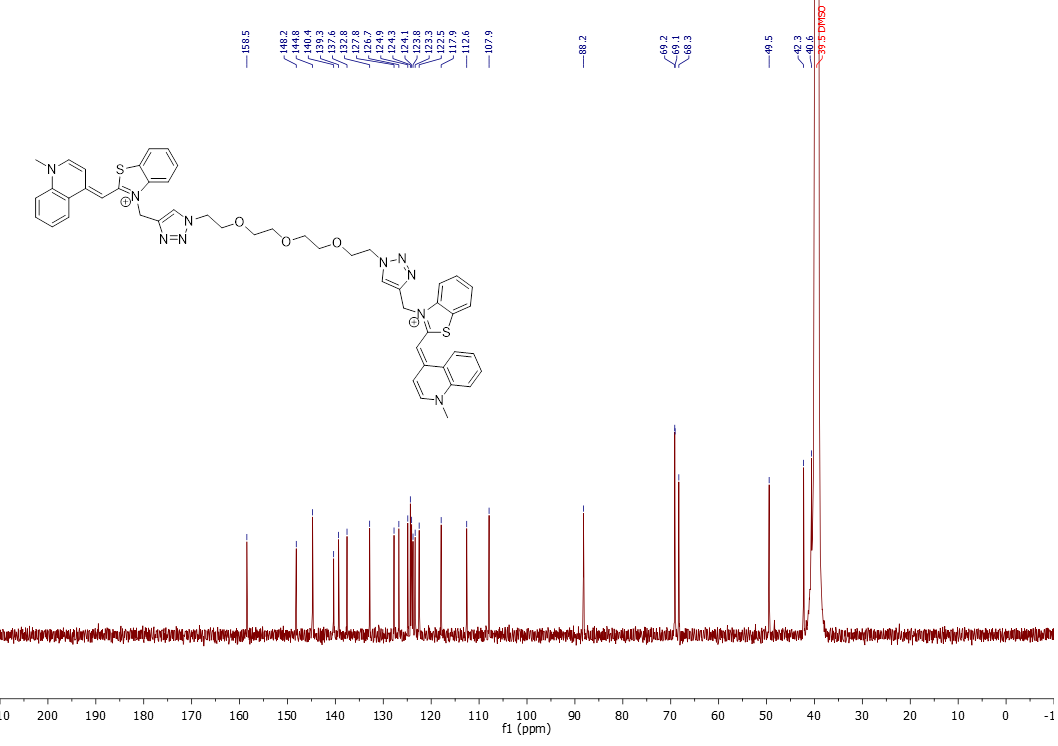

#### **Fig. S3. COSY spectrum of 3**

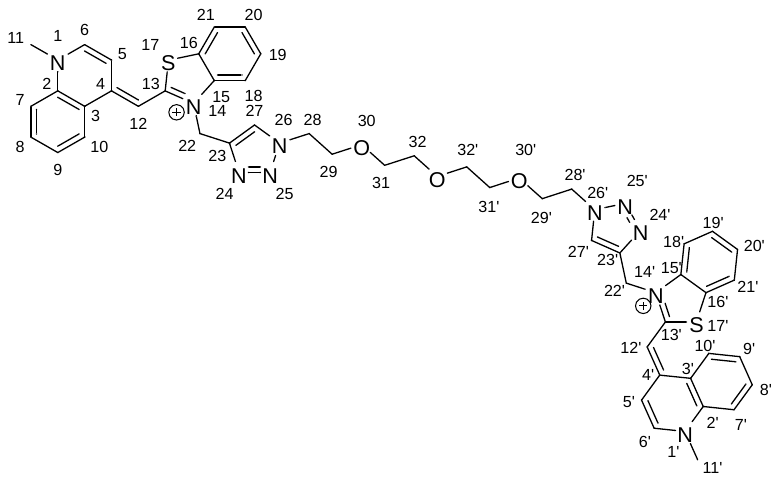

Aromatic region

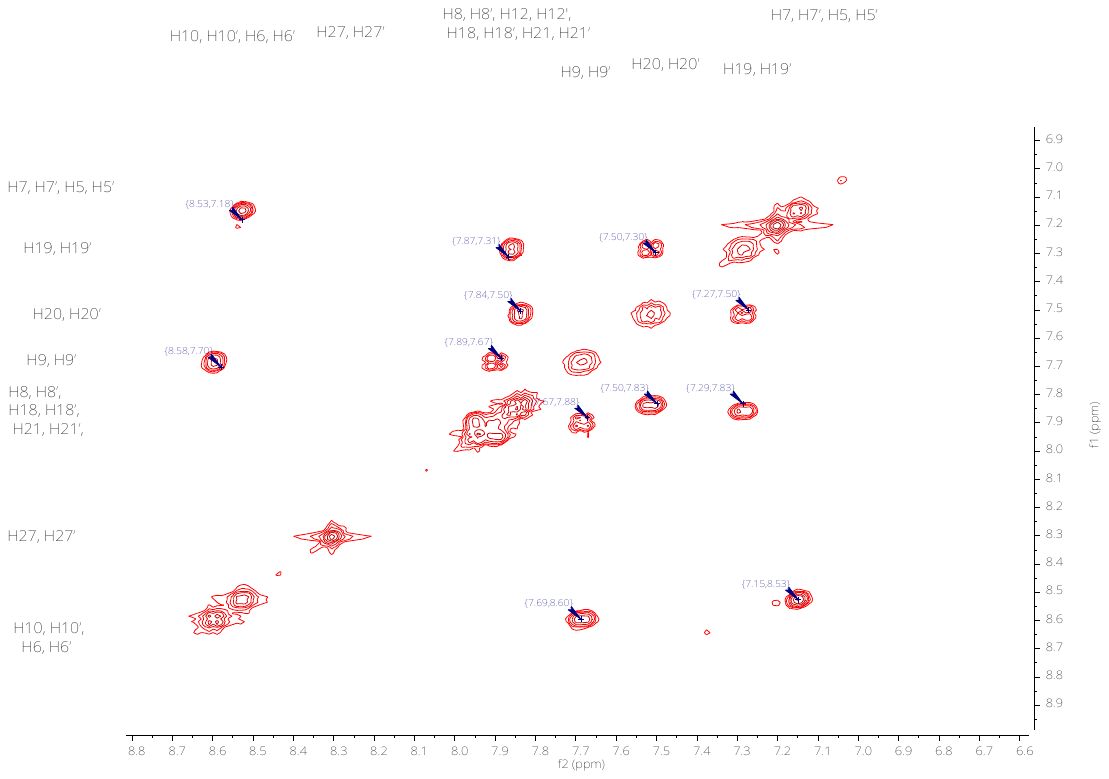

Aliphatic region

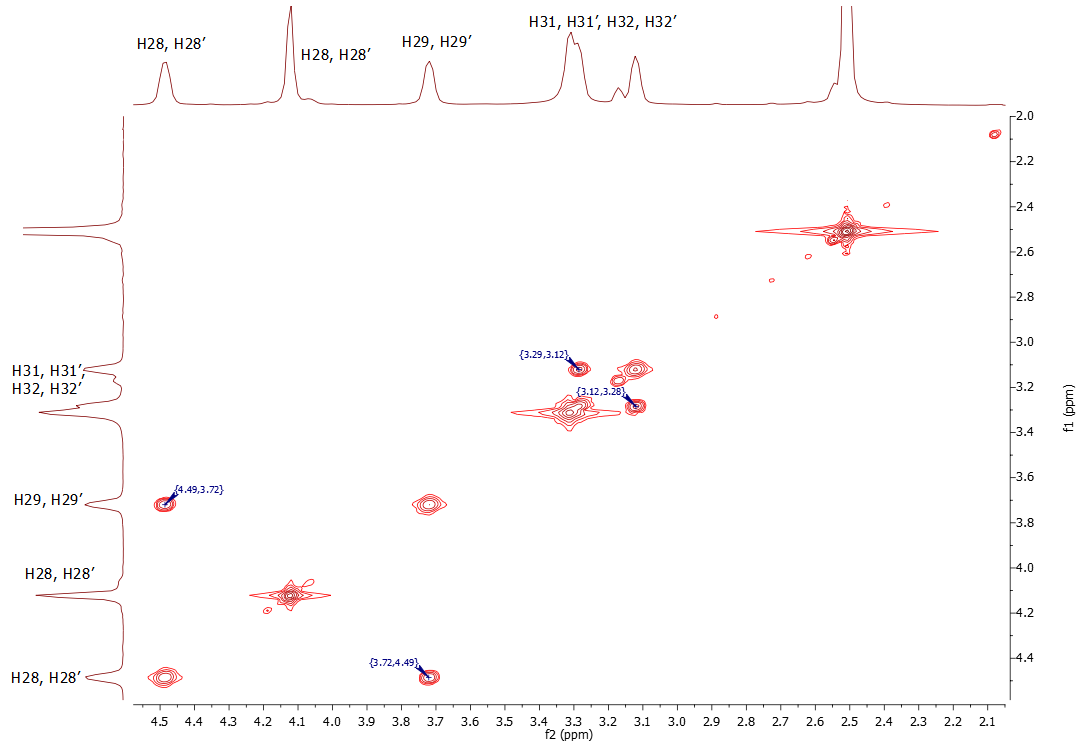

Full spectrum

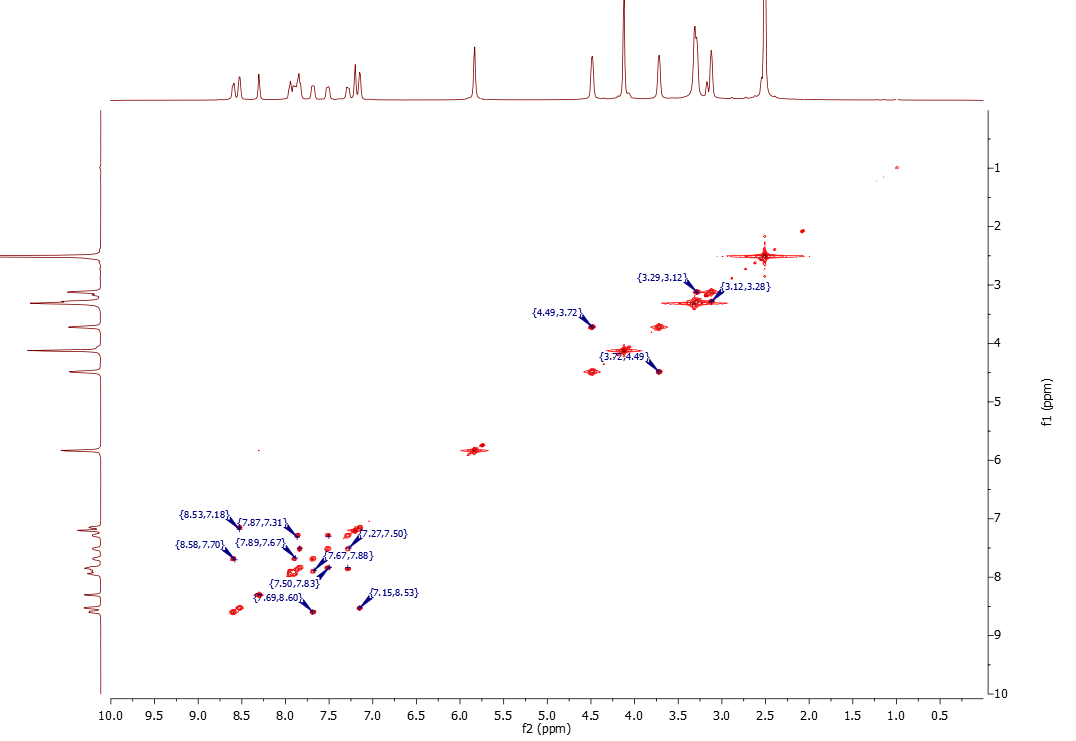

#### **Fig. S4. ^1^H spectrum of 4**

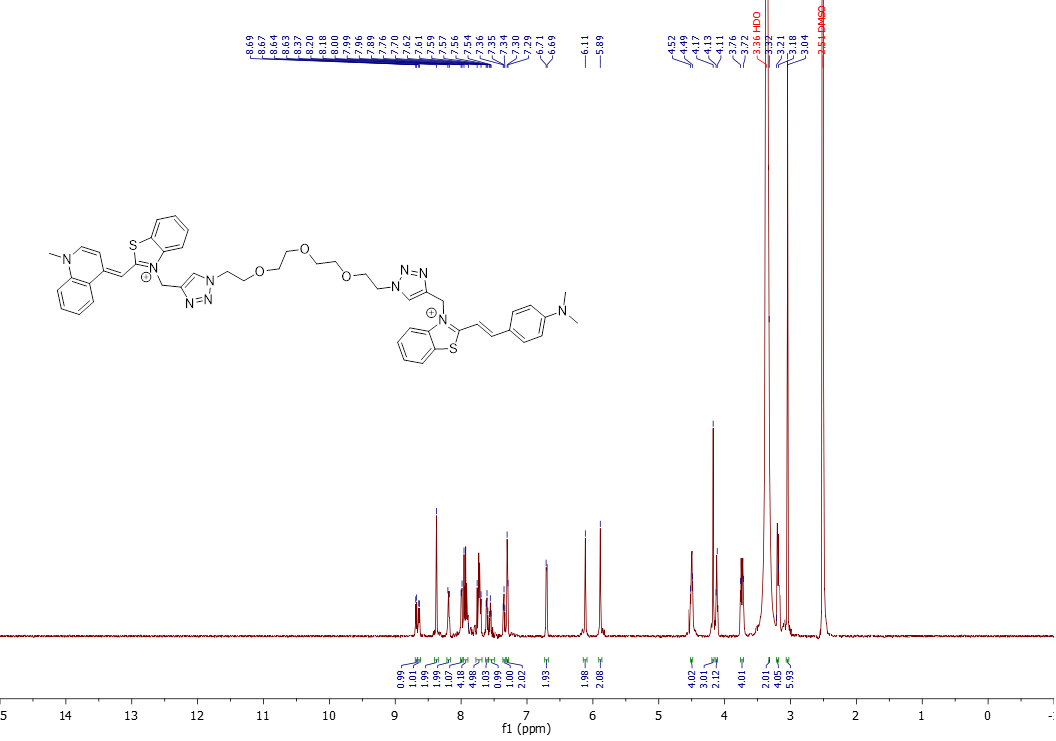

#### **Fig. S5. COSY spectrum of 4**

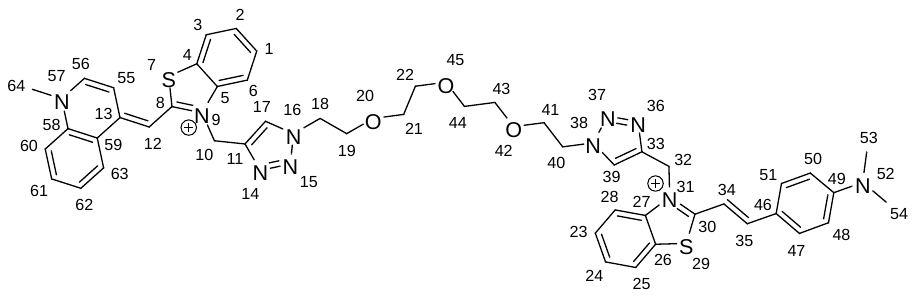

Aromatic region

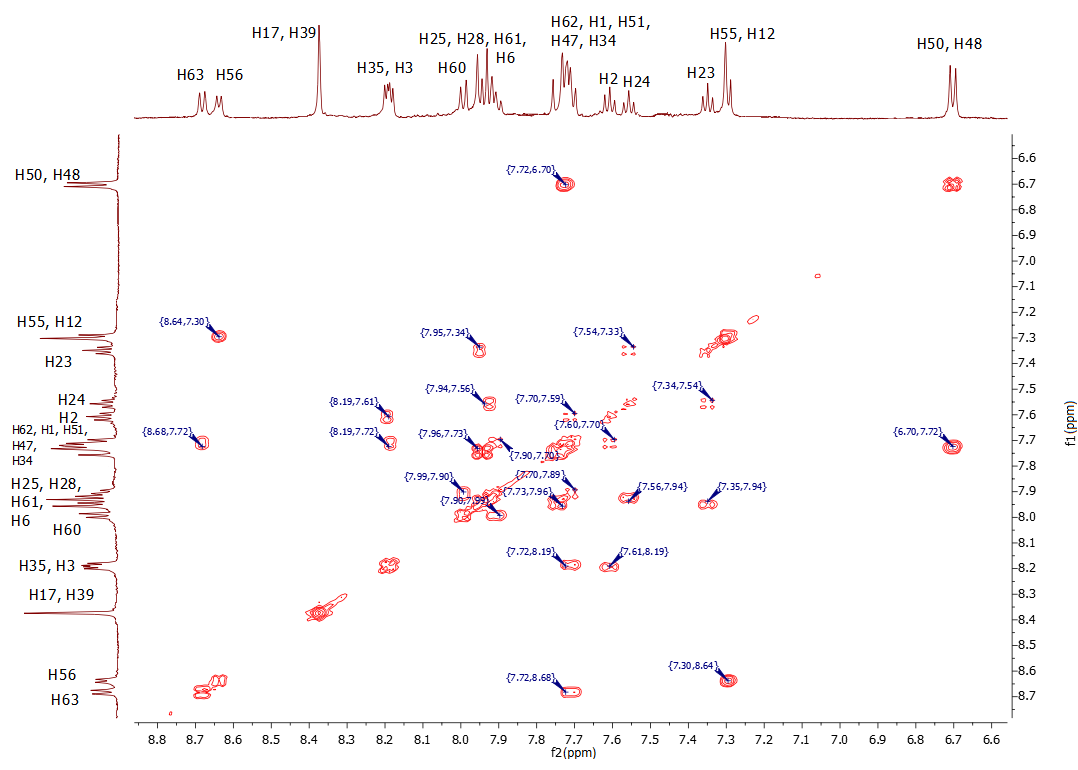

Aliphatic region

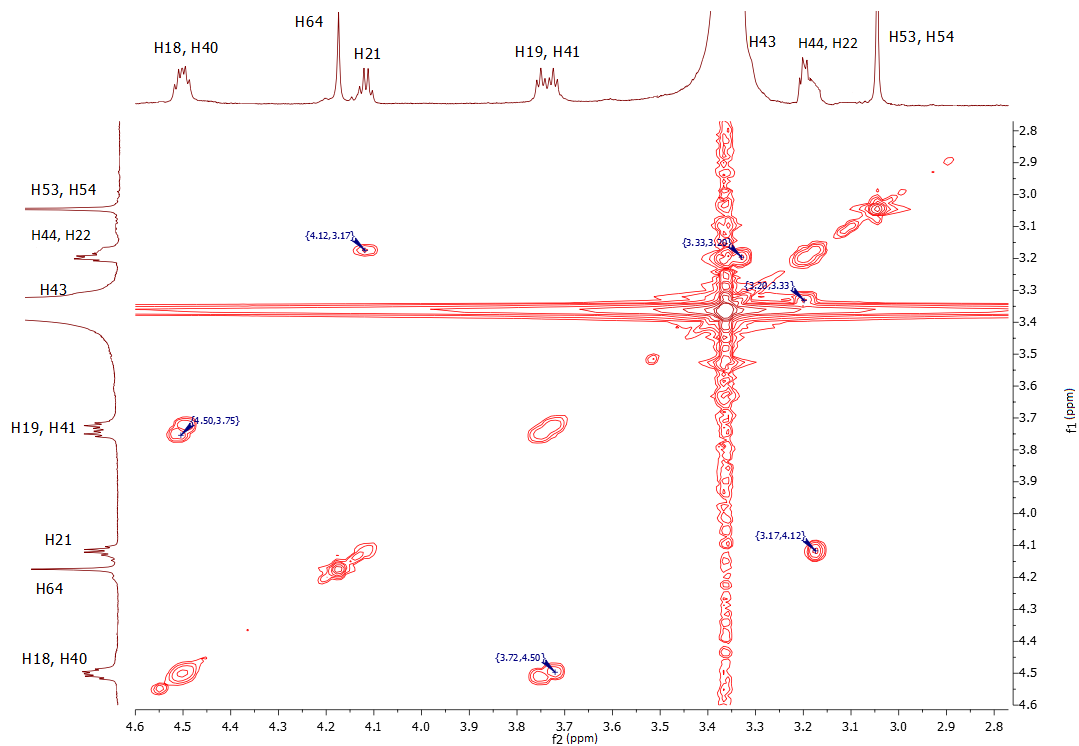

Full spectrum

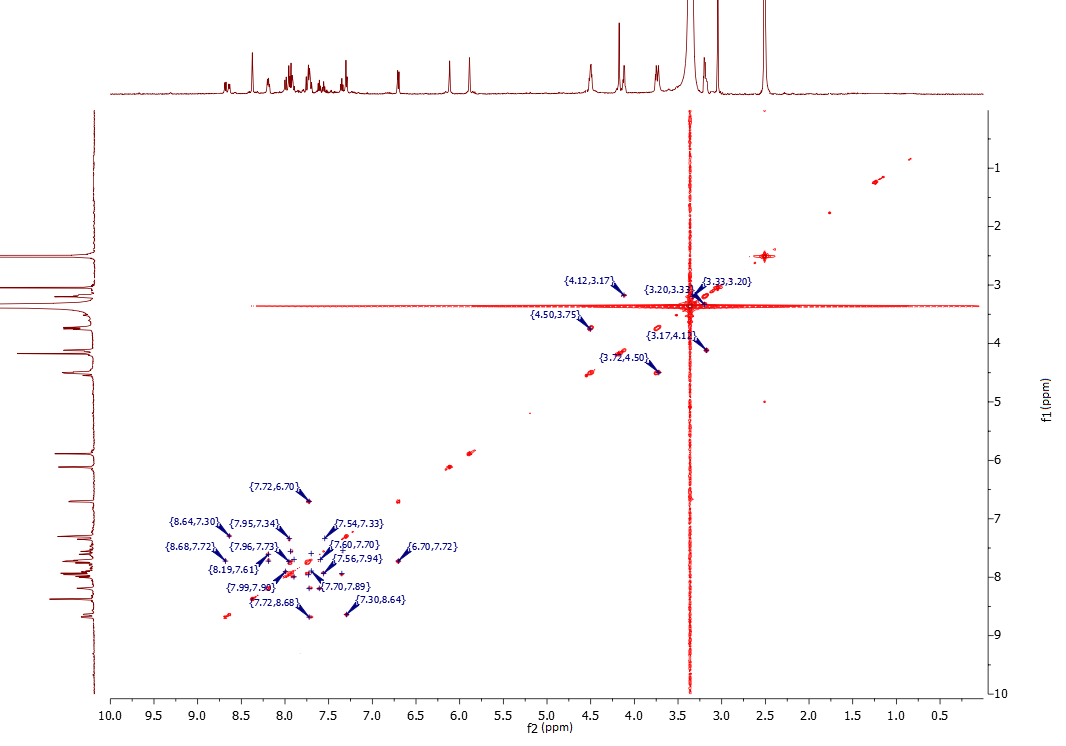

#### **Fig. S7. ^1^H spectrum of 6a**

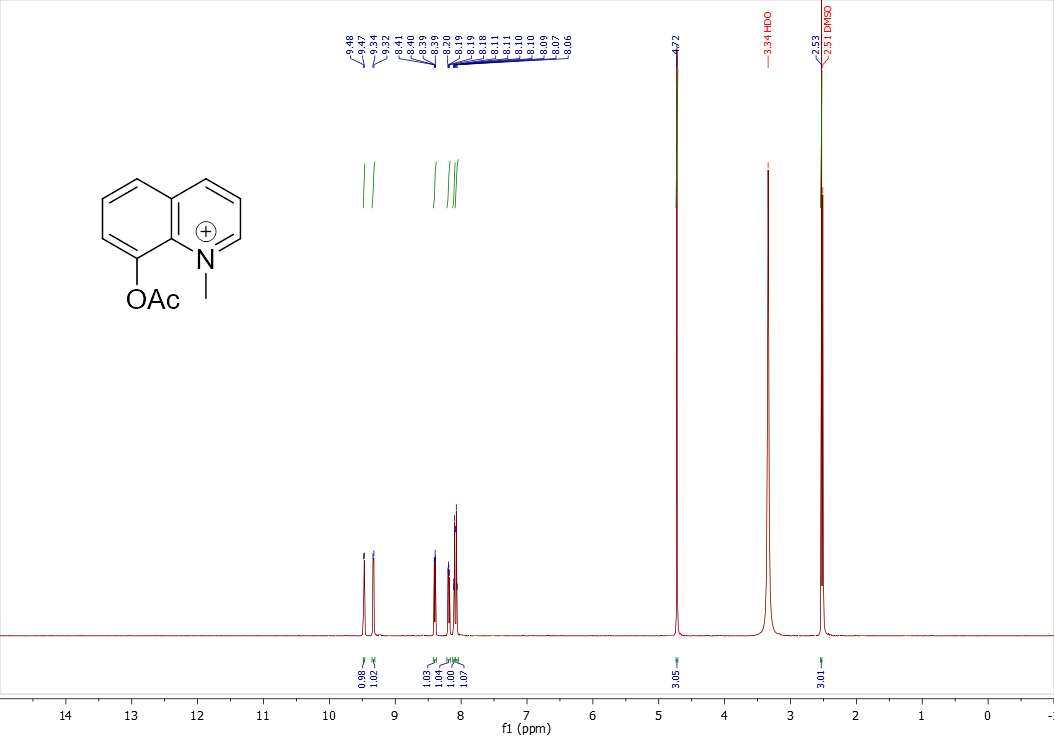

#### **Fig. S8. ^13^C spectrum of 6a**

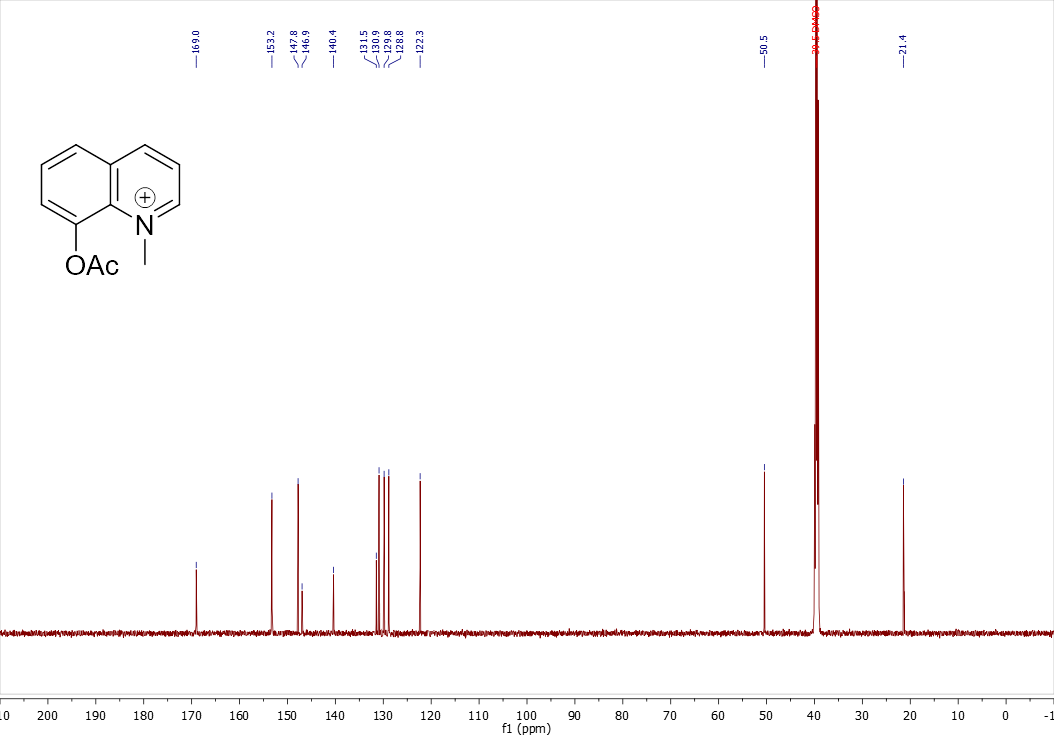

#### **Fig. S9. ^1^H spectrum of 7b**

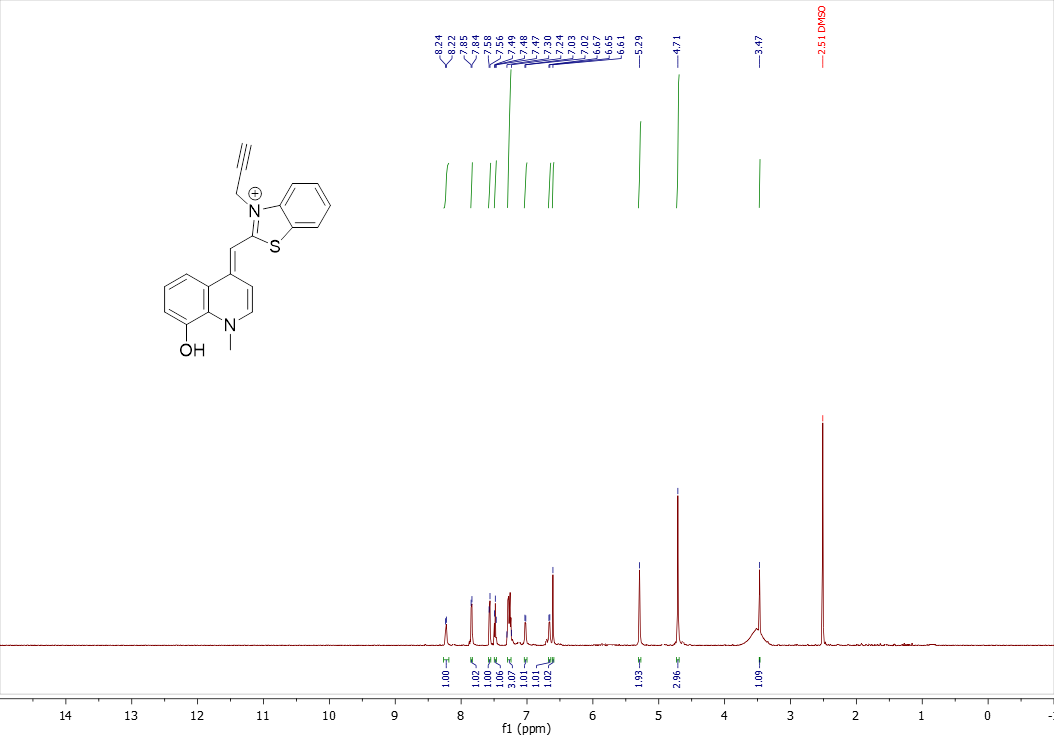

#### **Fig. S10. ^13^C spectrum of 7b**

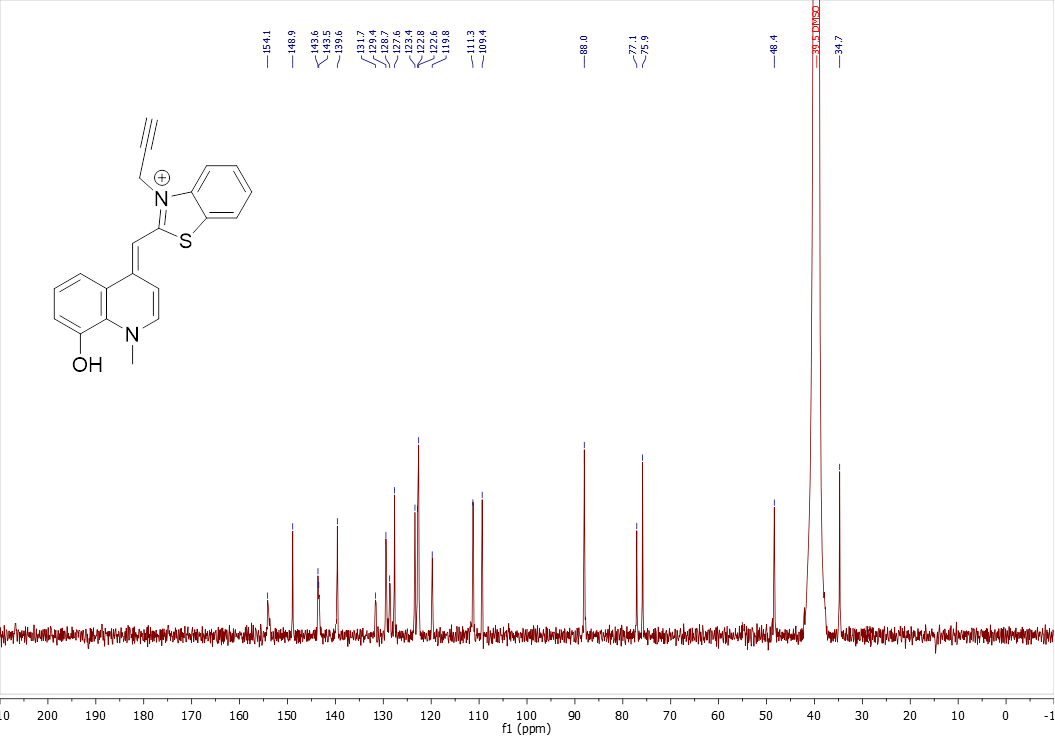

#### **Fig. S11. ^1^H spectrum of 7c**

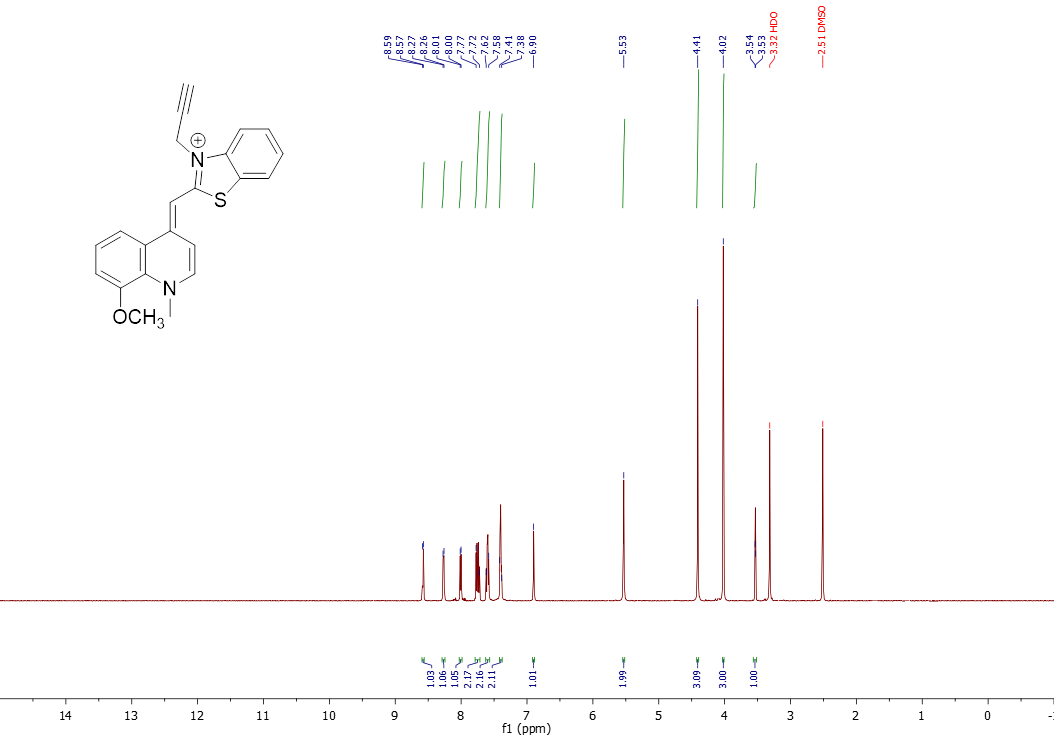

#### **Fig. S12. ^13^C spectrum of 7c**

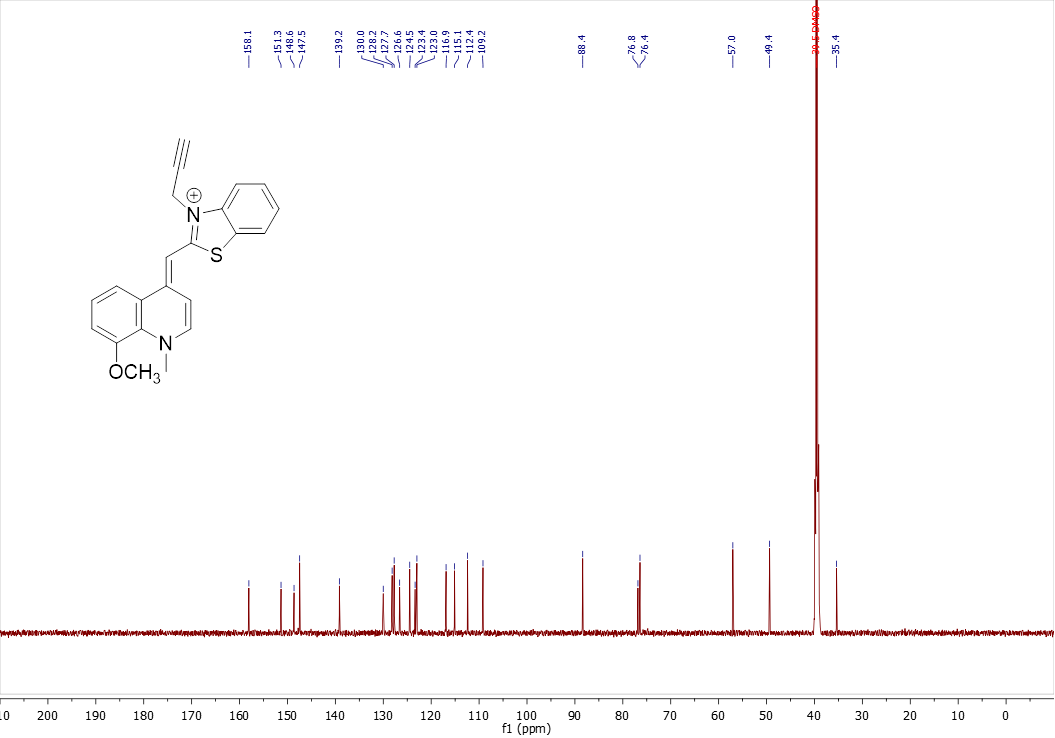

#### **Fig. S13. ^1^H spectrum of 9a**

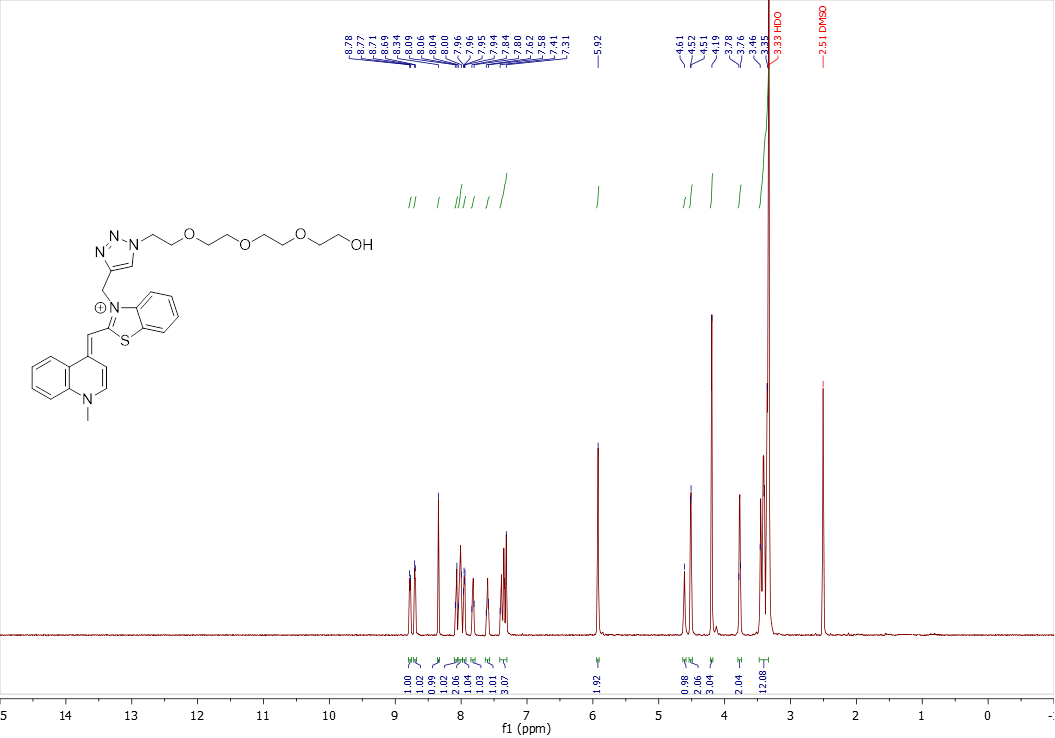

#### **Fig. S14. ^13^C spectrum of 9a**

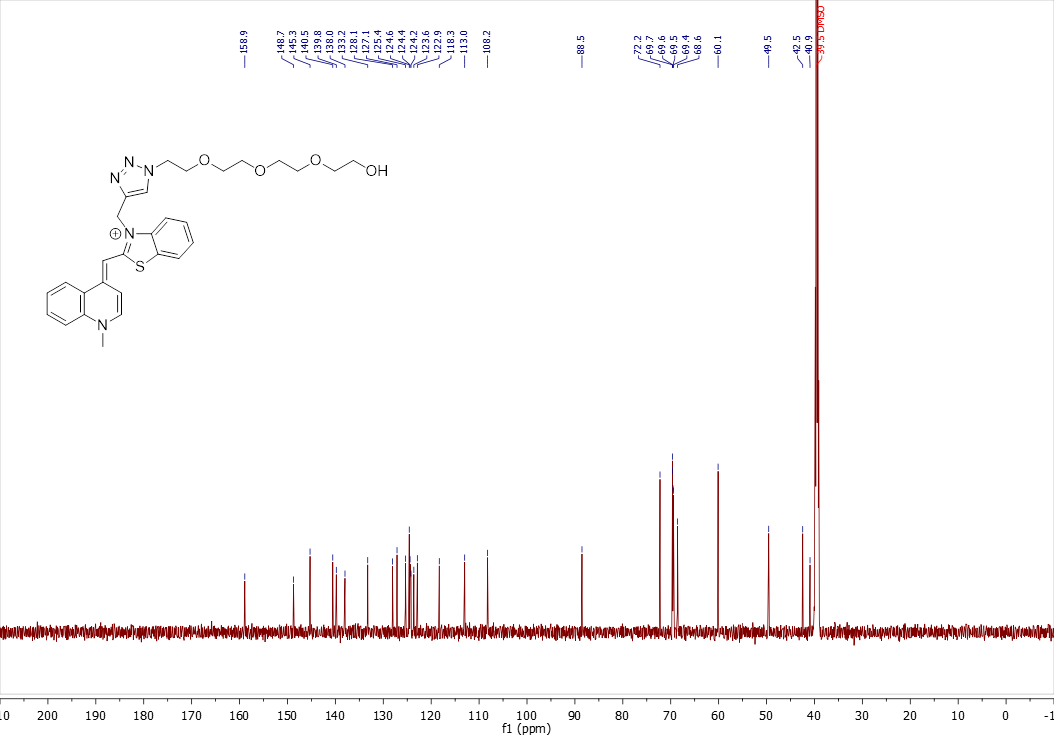

#### **Fig. S15. COSY spectrum of 9a**

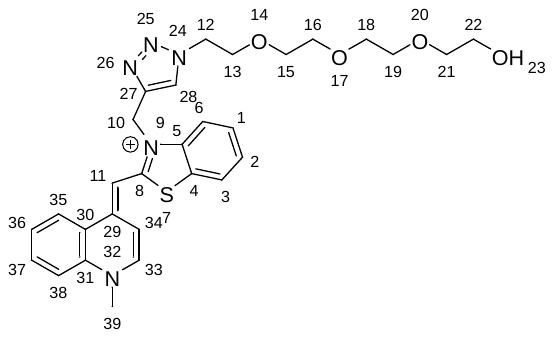

Aromatic region

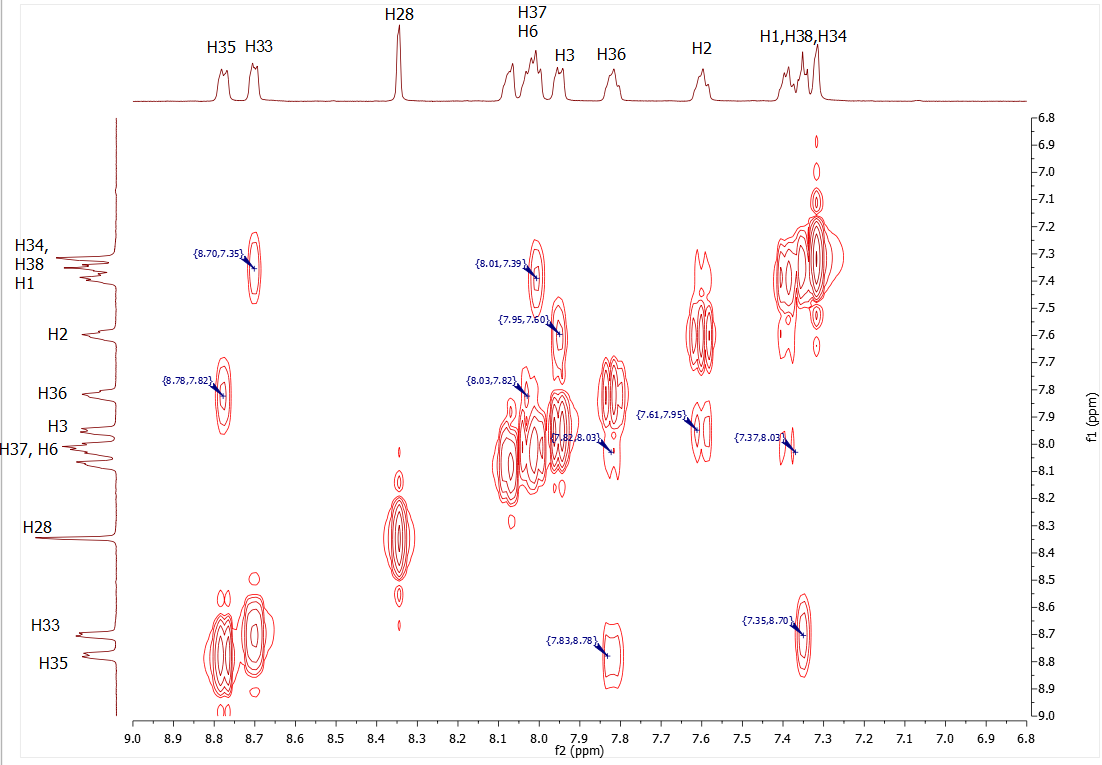

Aliphatic region

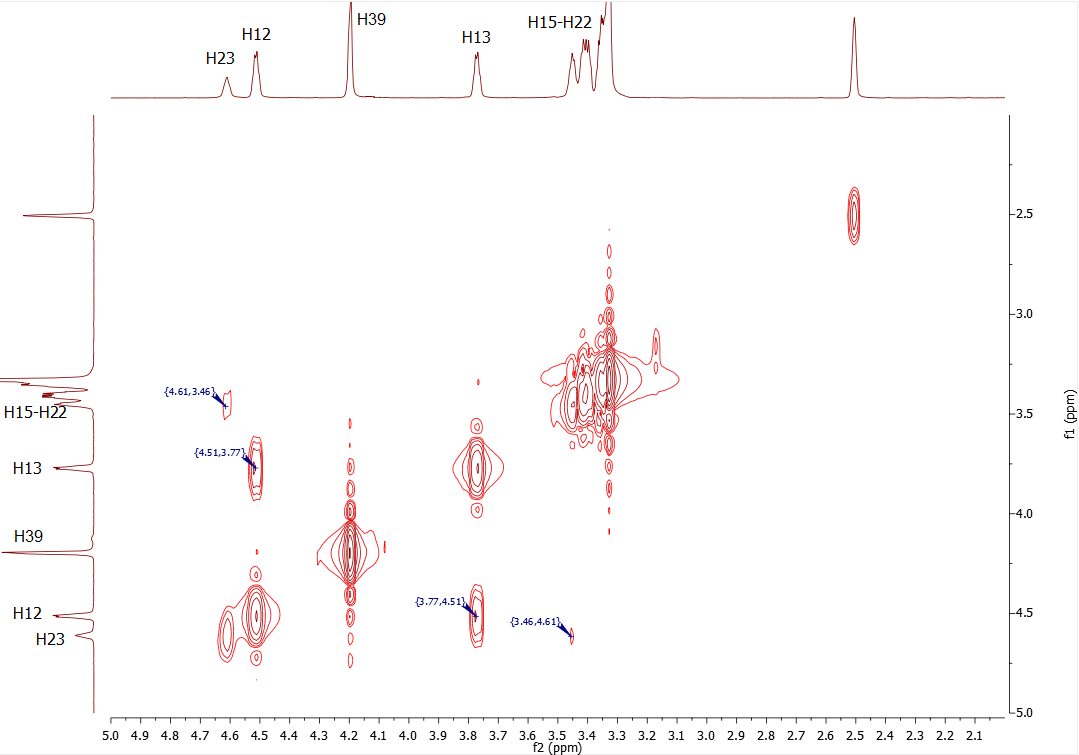

Full spectrum

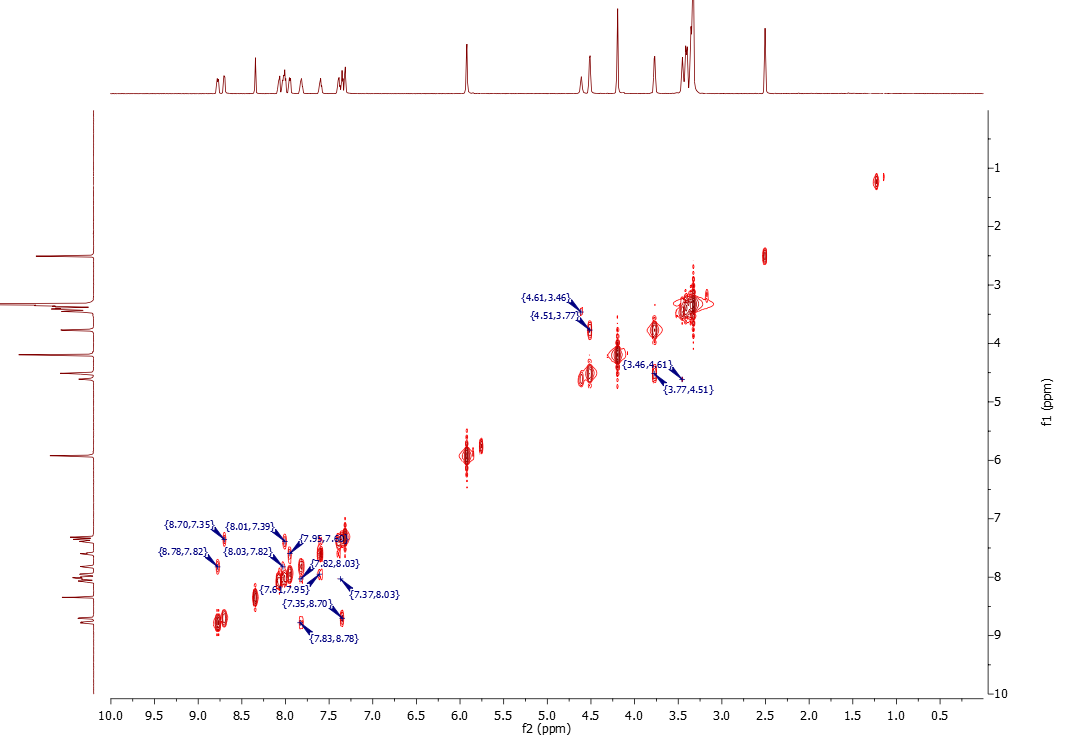

#### **Fig. S16. ^1^H spectrum of 9b**

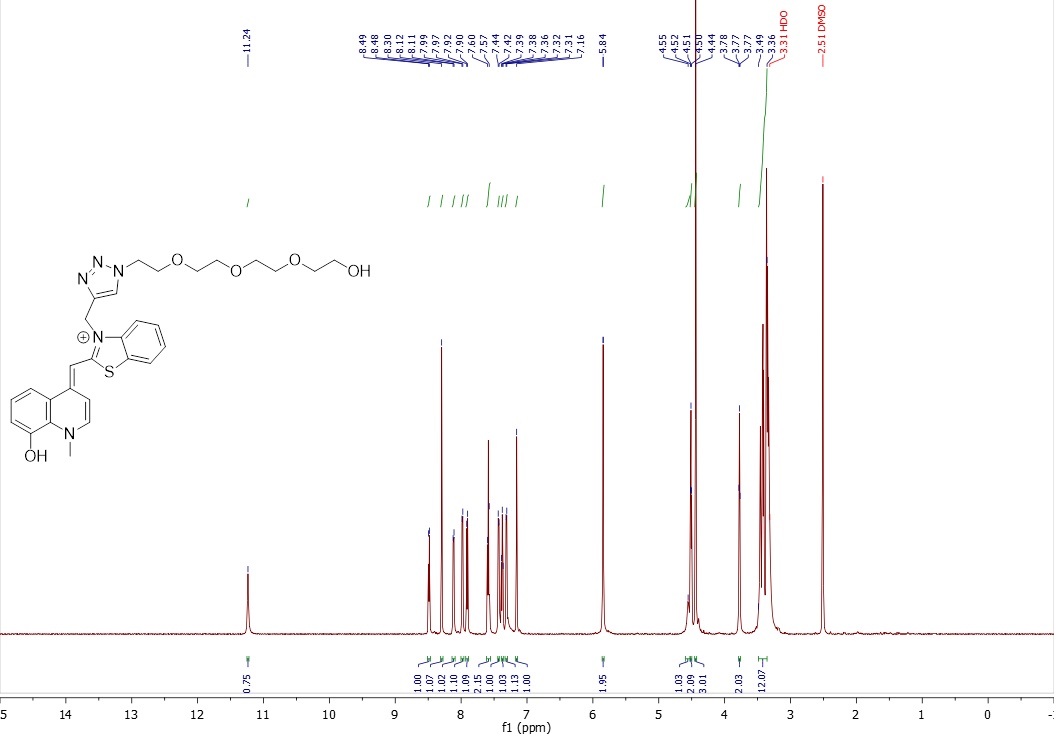

#### **Fig. S17. ^13^C spectrum of 9b**

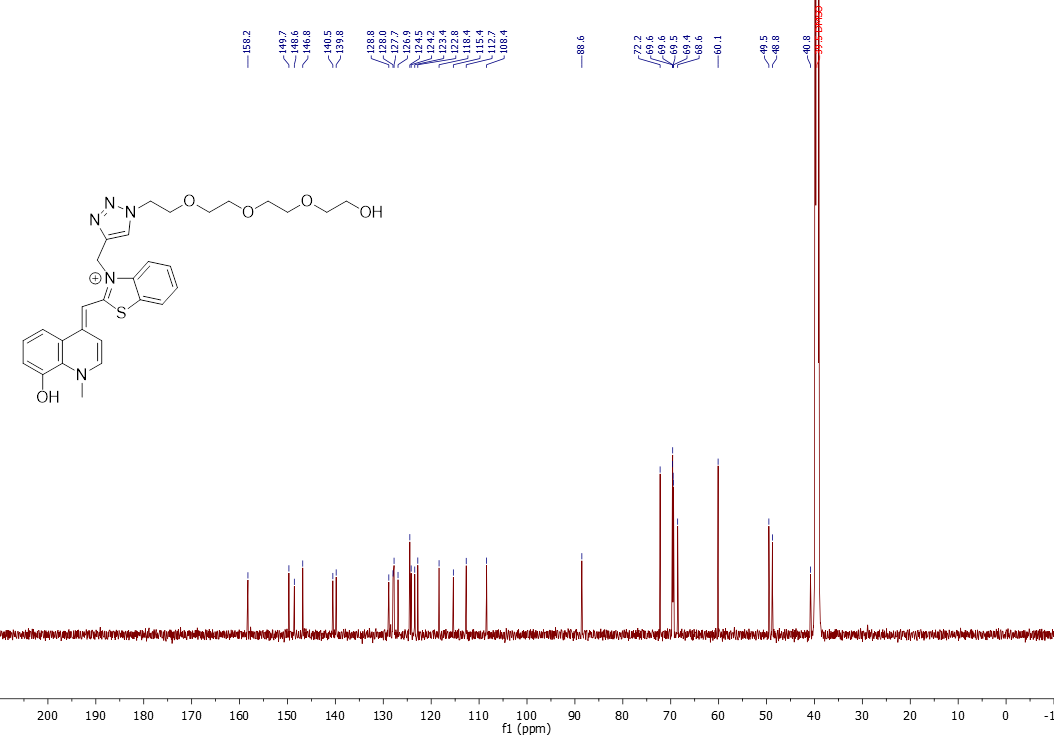

#### **Fig. S18. COSY spectrum of 9b**

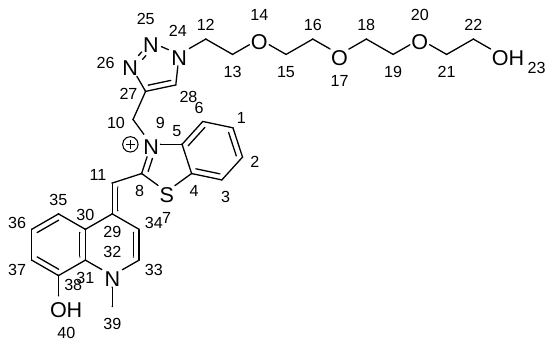

Aromatic region

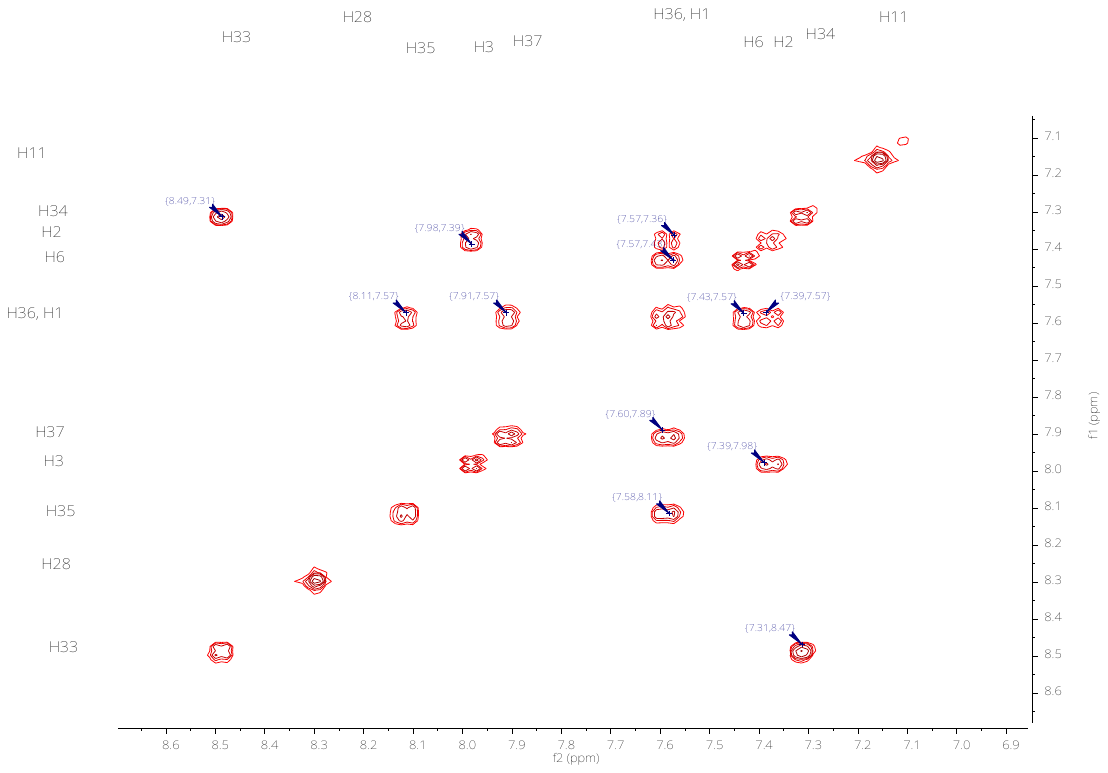

Aliphatic region

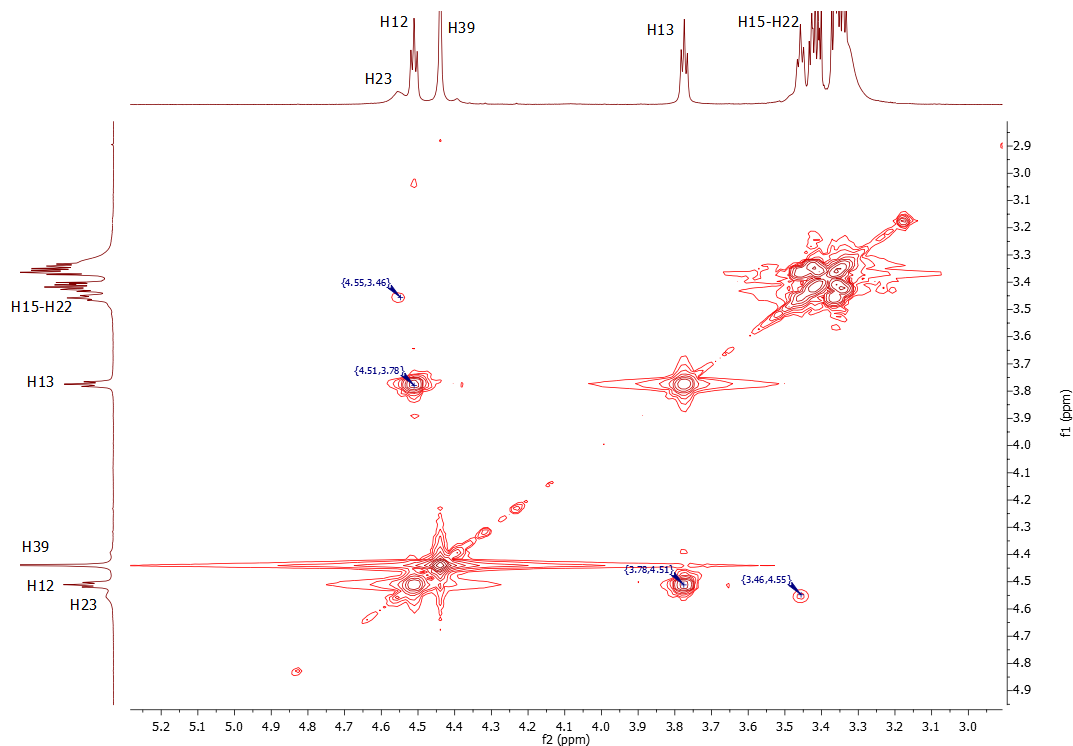

Full spectrum

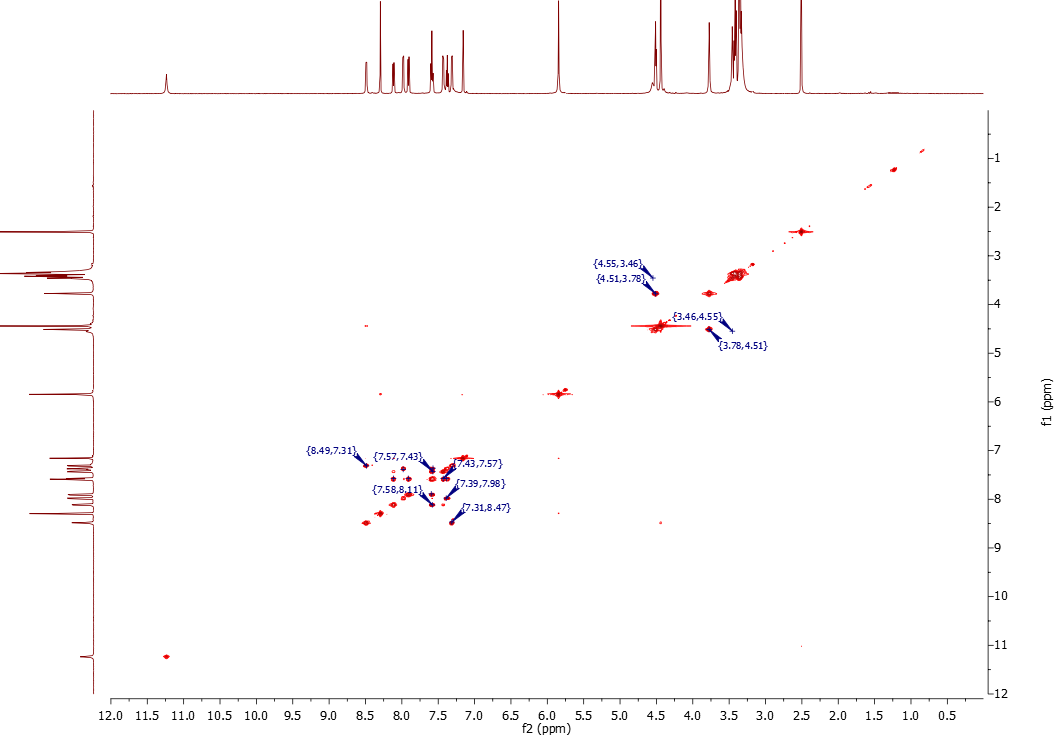

#### **Fig. S19. ^1^H spectrum of 9c**

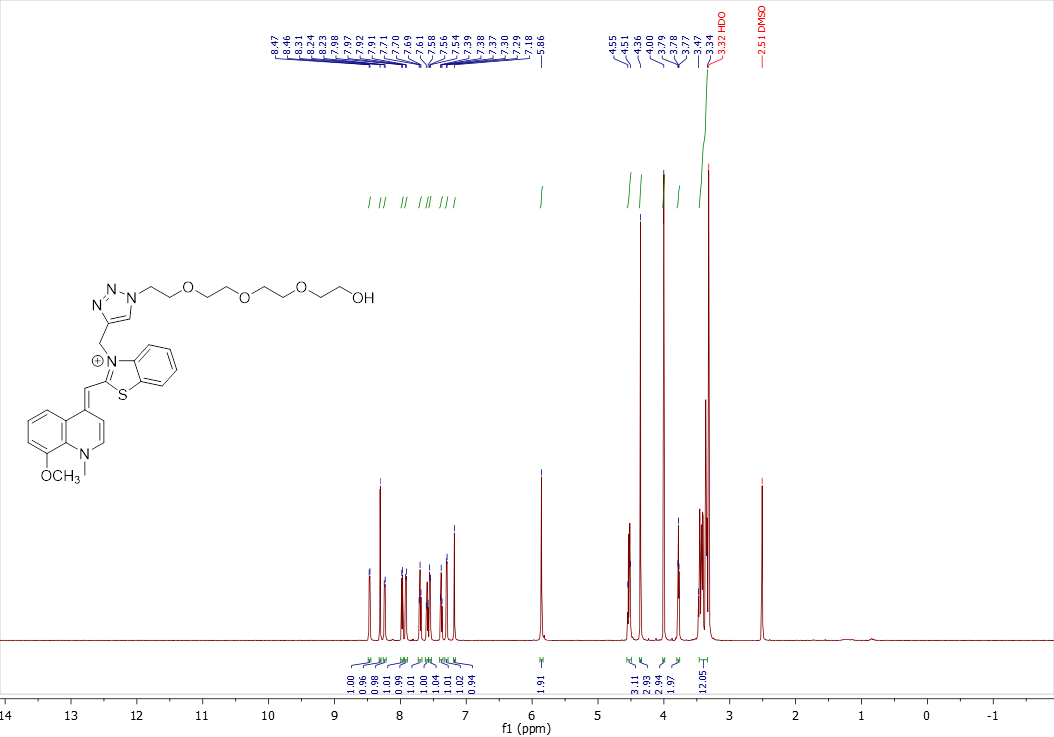

#### **Fig. S20. ^13^C spectrum of 9c**

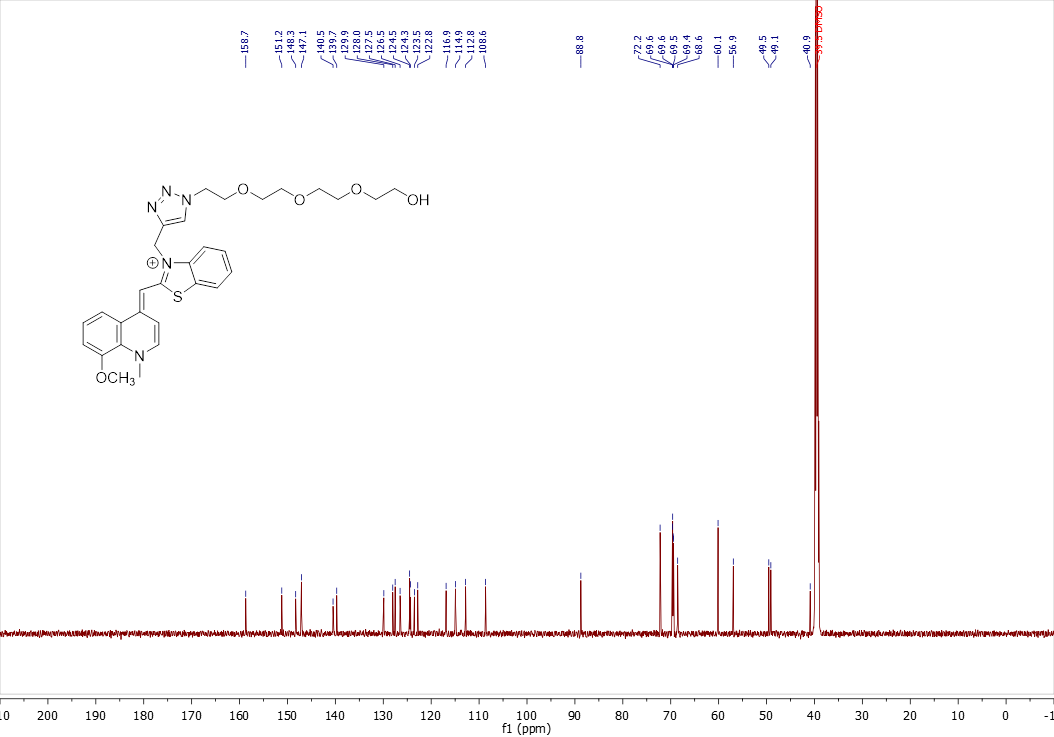

#### **Fig. S21. COSY spectrum of 9c**

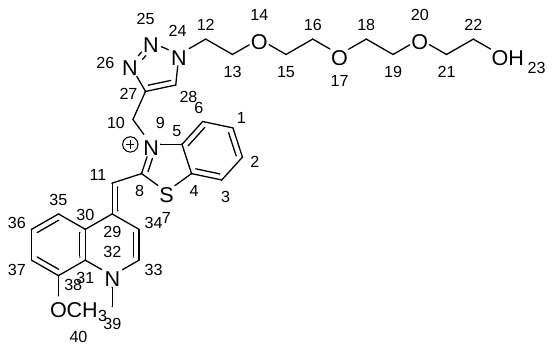

Aromatic region

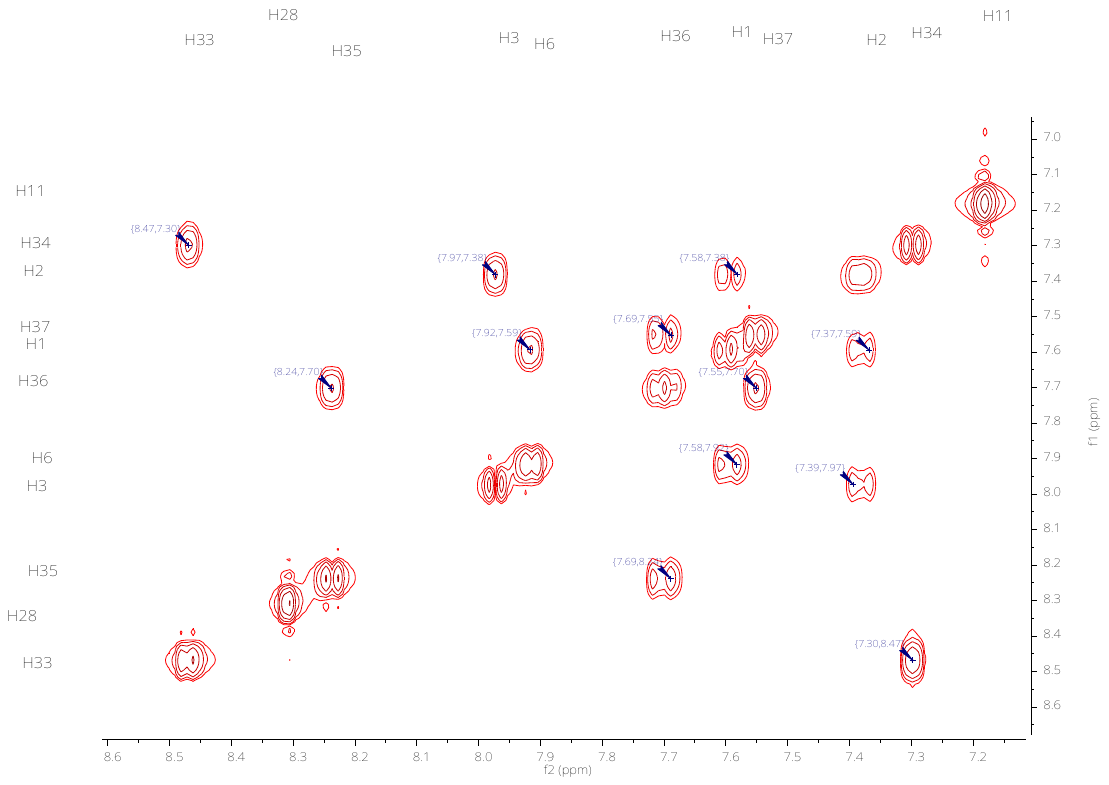

Aliphatic region

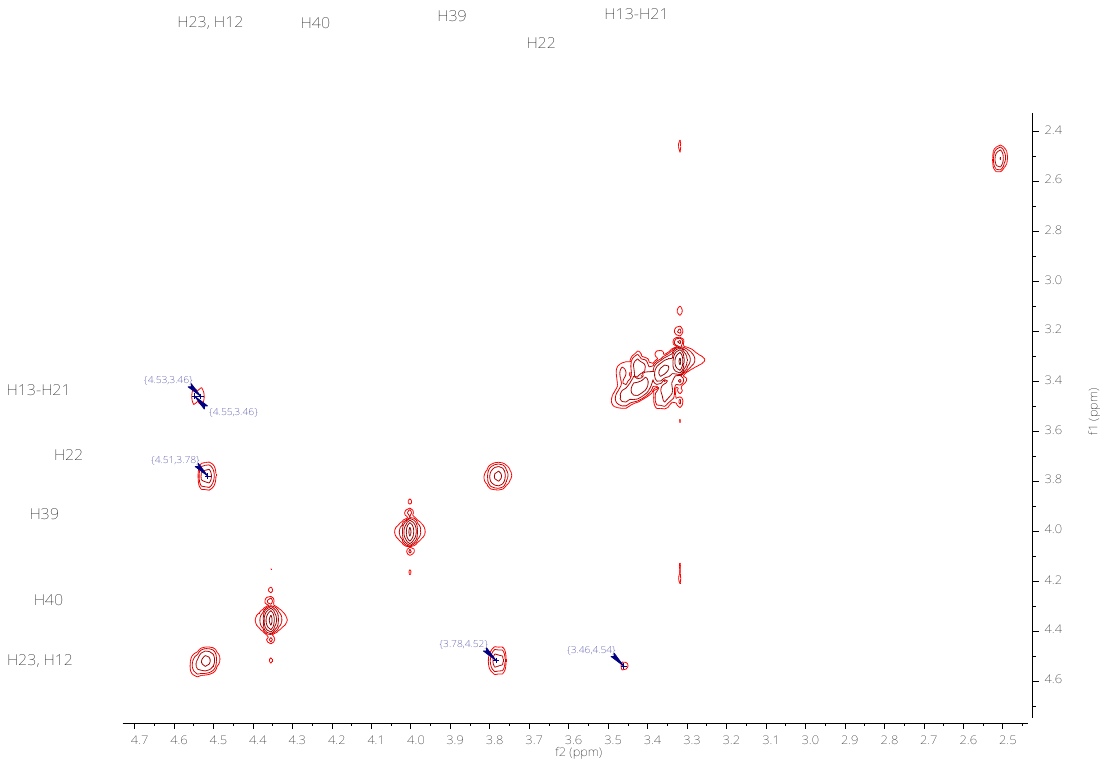

Full spectrum

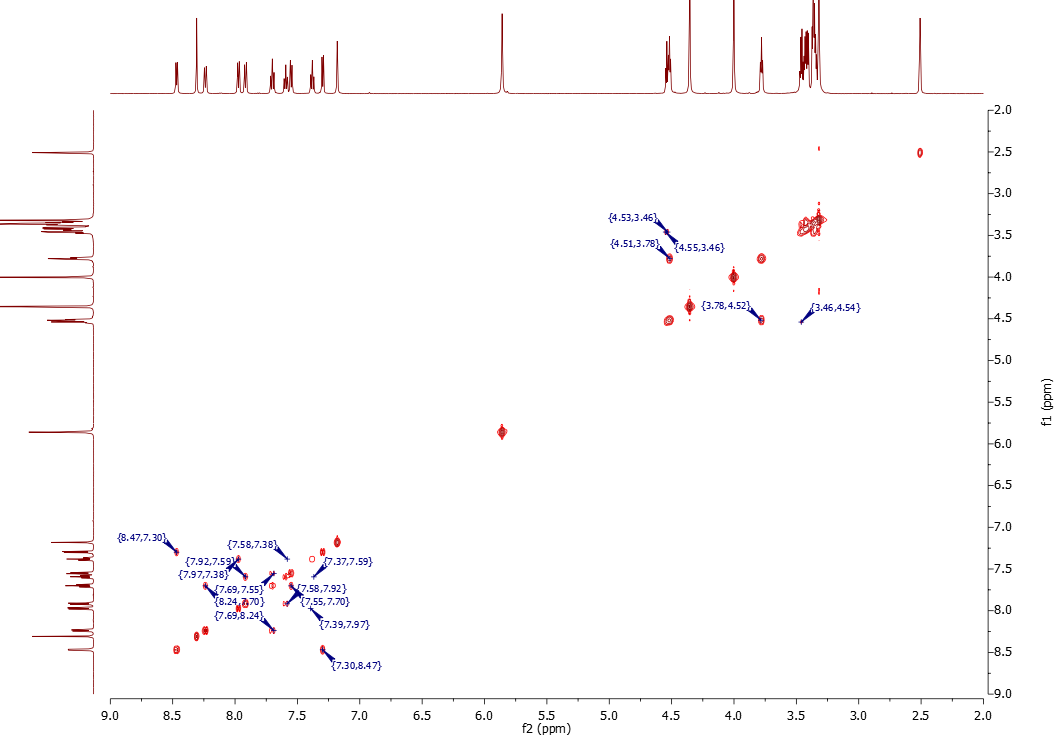
